# Cellular basis of accelerated whole-tooth regeneration

**DOI:** 10.64898/2026.01.06.697137

**Authors:** Talha Mubeen, Haowen He, George W. Gruenhagen, Anoushka Satoskar, Jeffrey T. Streelman

## Abstract

Teeth are ectodermal organs that have, throughout their long evolutionary history, retained the capacity for full regeneration and replacement, even in adult stages. Yet, because most mammals (e.g., humans, mice) lack lifelong dental replacement, we do not fully understand its tempo and mode, and we do not have a clear picture of the cell populations and signals that contribute to the process. Here, we used cichlid fishes from Lake Malawi, species that differ in tooth formula (tooth shape and number) but share one-for-one tooth replacement, to (i) explore the tempo of dental replacement after plucking and then (ii) identify the cell populations, gene expression signatures, and interactions between cell populations that change in this plucking paradigm.

We observed that cichlid species with divergent dentitions accelerated tooth replacement >3x on the plucked half of the jaw. Then, we used single-nucleus RNA-seq to profile cellular and molecular changes across the first week of post-plucking tooth replacement. This approach allowed us to infer cellular trajectories in dental epithelium and mesenchyme that underlie tooth regeneration. We identifed distinct gene expression profiles and cellular interactions across four time points of accelerated tooth replacement, with divergent involvement of epithelial, mesenchymal and immune cell types. Diferential signaling of Collagen, BMP, MMP, Semaphorin and Slit-Robo pathways was evident after plucking and highlights temporally-sequenced roles of immune response, odontogenesis, vascularization and nerve pathfinding as teeth are constructed anew. Overall, this study provides insight into the trajectory of cellular interactions accompanying whole-tooth replacement and offers a comparative foundation for understanding dental regeneration in vertebrates.

## INTRODUCTION

Most teeth in the half-billion year evolutionary history of dentitions exhibited the capacity for replacement. Despite the deep conservation of molecular programs used to build the unit tooth^1^, dental pattern and mode of regeneration differ dramatically across vertebrates. Birds, for instance, lack teeth entirely; most fishes have multiple rows of replacing dentitions on two jaws, while mammals possess teeth in a single row at the oral jaw margin with limited potential for replacement. Implications of variance in tooth replacement across vertebrates are manifold. Because mice don’t replace their teeth even once, there is no traditional laboratory model for tooth replacement. In humans, where tooth replacement occurs once in a lifetime, 23% of people over the age of 60 lack teeth altogether^2^.

There is a voluminous literature on tooth development^3^, and as with other organs, speculation that developmental signals also govern regeneration^4^. Vertebrates with lifelong tooth replacement tend to possess a successional dental lamina, a band of epithelium linking the functional to replacement tooth. The successional lamina (SL) holds label-retaining cells, and expresses stem cell markers as well as markers of dental competence in lizards, alligators, sharks and fishes^5–11^. Notably, stickleback fishes replace their teeth and lack a successional lamina, but they express conserved markers in the vicinity of tooth replacement^12^.

Tooth development proceeds through stages of reciprocal signaling between dental epithelium and neural crest derived ectomesenchyme. Tissue renewal of the epithelium is well studied in the mouse incisor^13–15^, where a conveyor-belt of Sox2+ epithelial stem cells (ESCs), and their derivatives, align along the labial surface to replace enamel and other epithelial lineages, lost from constant gnawing. The situation is more complicated in mesenchyme of mouse incisors. Classic lineage-tracing studies identified multiple sources of mesenchymal stem cells (MSCs), evidence of transdifferentiation after injury and perhaps different mechanisms during homeostasis versus repair^16–18^. Even though both epithelium and mesenchyme are required to make a tooth, murine dental ESCs and MSCs have been studied separately as exemplars of epithelial and mesenchymal regeneration, respectively.

Reductionist approaches like single-cell or single-nucleus RNA-sequencing (scRNA-seq; snRNA-seq) have rekindled integrative thinking about whole-tooth biology, with the aim of assembling a parts list of dental cell types and identifying how cell populations work together to build or to repair a dentition^19^. Studies in mouse and human teeth catalogued and compared dental cell types^20–23^. Computational analyses uncovered lineage dynamics of cell states during ameloblast and odontoblast differentiation, during both homeostasis and injury repair. Finally, work has revealed a deep conservation of cell type and cell type gene expression across vertebrate teeth, despite hundreds of millions of years of evolution^24^.

Vetebrate teeth that undergo lifelong replacement share homologous cell types with non-replacing mammalian dentitions^24^, and analogous processes of cyclic renewal with other vertebrate ectodermal organ systems like hair and feathers^25–27^. These systems share features like (i) stem cell niches in epithelium and mesenchyme sufficient to build organs anew, (ii) a timing mechanism to govern and order the process of replacement and (iii) contingency to accelerate replacement when organs are damaged. Over the past two decades, we developed the genomic resources^28^ and molecular approaches to study the coupled processes of dental patterning and tooth regeneration in Lake Malawi cichlids, a group of fishes known for rapid and rampant evolution of phenotypes and behaviors. These species share similar genomes, but exhibit dramatic differences in tooth shape, tooth number and tooth spacing^29^. The tooth pattern (spacing/size of teeth) is set early in development^30^ (first few weeks of life) and then in juvenile and adult stages, each tooth is replaced with shape fidelity every 30-40 days^10^. Cichlid teeth are co-patterned with taste buds on oral and pharyngeal jaws by competing Hh, Bmp and Wnt signals^31^ and one-for-one tooth replacement is galvanized by multiple niches that house label-retaining cells and express ESC and MSC markers^11^. In this report, we use the Malawi cichlid system to explore the tempo and mode of whole-tooth replacement by first establishing a tooth plucking paradigm across species with divergent dental formulae. Replacement was accelerated on the plucked side of the jaw, so we then employed snRNA-seq in this paradigm to identify cell types and molecular signals that coordinate the process.

## RESULTS

### Plucking accelerates tooth replacement on the plucked side

We aimed to manipulate and quantify tooth replacement in the Malawi cichlid system. To do so, we coupled whole-tooth plucking, on one half of the jaw, with a pulse-chase method that differentiates old from new teeth, after a 15-day recovery period^32^ (Figure 1A-B; Supplemental Figure 2). In this set-up, each individual is a replicate, with jaws split by control vs plucked conditions. We established this paradigm in three rock-dwelling Malawi cichlids with divergent dentitions: (1) *Cynotilapia afra* (CA), which exhibit 2-3 rows of widely-spaced, large conical unicuspid teeth (Supplementary Figure 1A-C); (2) *Metriaclima estherae* (MZ Red) which possess an outer row of large bicuspid teeth followed by multiple rows of smaller tightly packed tricuspids in posterior rows (Supplementary Figure 1D-F); and (3) *Petrotilapia chitimba* Thick Bar (PT) whose jaw is lined with multiple rows of tightly packed curved tricuspid teeth (Supplementary Figure 1G-I). Tooth plucking mimics, to some degree, natural tooth loss among males, due to frontal display and assault in aggressive rock-dwelling species^33^.

**Figure 1.**
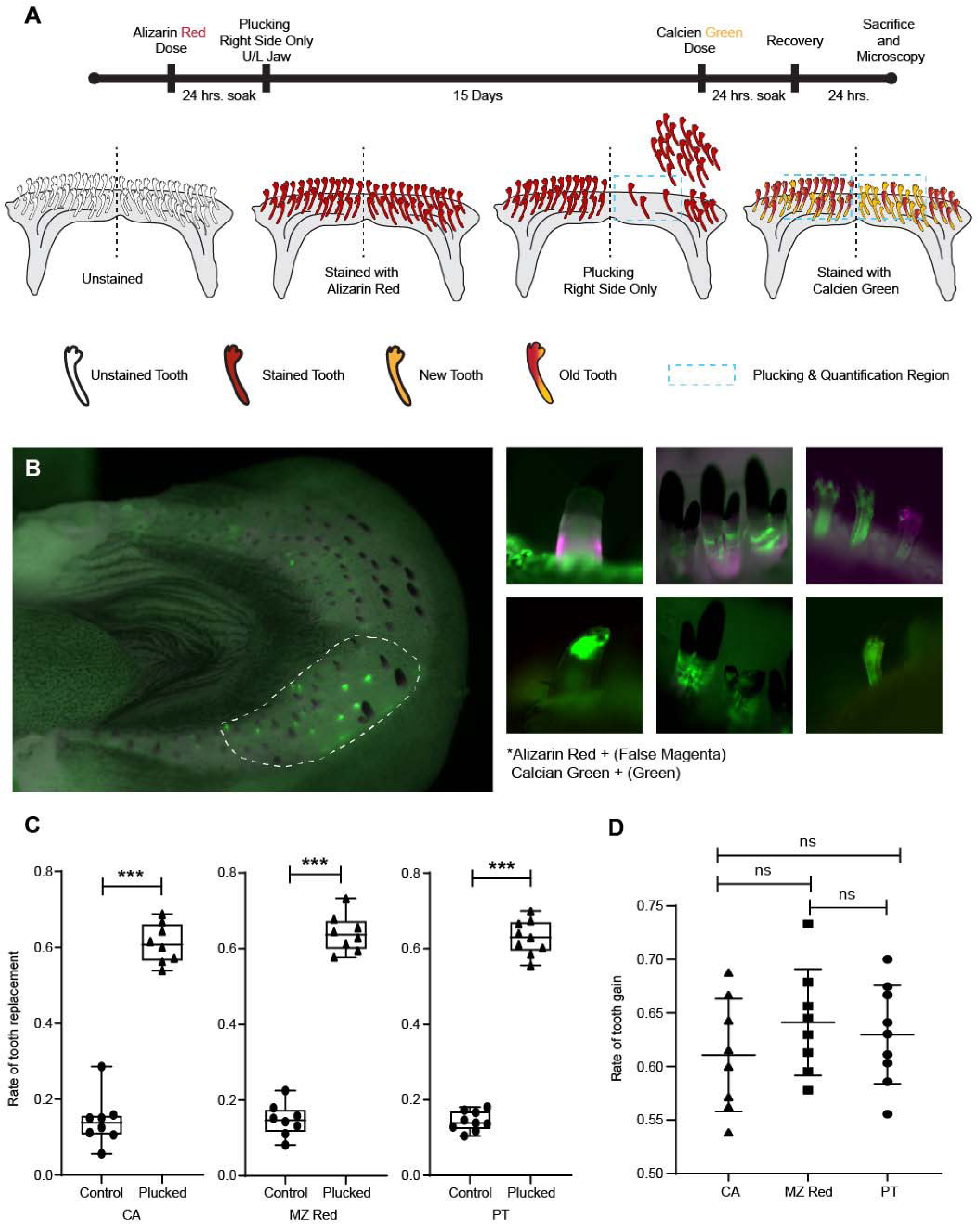
Tooth replacement is accelerated after plucking in cichlid species with divergent dentitions. A. Schematic of the modified pulse-chase method using Alizarin and Calcein to quantify tooth replacement. B. Representative fluorescence images used to classify teeth by dye incorporation. Teeth positive for both Alizarin and Calcein were scored as pre-existing old teeth, whereas teeth positive for Calcein only were scored as newly formed teeth. Left, whole jaw from MZ Red, with teeth plucked on the right side and the left side serving as the control; the dotted outline marks the region used for plucking and quantification (Supplemental Figure 2B). Right, representative individually classified teeth (Alizarin, magenta; Calcein, green), with the top row showing pre-existing old teeth positive for doth dyes and the bottom row showing newly formed teeth positive only for Calcein. C. Pulse-chase reveals an elevated rate of tooth replacement on the plucked side in CA, MZ Red and PT. Tooth gain is shown as the normalized replacement rate (new teeth/total teeth) \*\*\**p*-value < 0.0001, paired Student’s t-test; CA (n = 8), MZ Red (n = 8) and PT (n = 9). D. Rate of tooth gain is consistent across species after plucking. NS, *p*-value > 0.05, one-way ANOVA with Tukey-Kramer post-hoc test; CA (n = 8), MZ Red (n = 8), PT (n = 9).

We note here key aspects of tooth replacement in Malawi cichlids^10,11,29,31^. In this system, tooth replacement occurs in a one-for-one manner with every other tooth in synch (such that neighboring teeth are typically not undergoing replacement at once). When a functional tooth is shed, the replacement completes development and erupts, with now the next tooth bud in the family poised beneath it. Notably for our case then, experimental and control sides of the jaw will have approximately the same number of replacement teeth at the onset of plucking (as illustrated in Figure 2A), but those on the plucked side are now simultaneously mobilized to develop, grow and erupt. Depending on the stage of the tooth in the ‘ready’ position, replacement tooth buds must also recruit nerves and blood vessels. Tooth replacement continues on the ‘control’ side of the jaw. Because of pronounced natural variation in tooth size, height and eruption stage, complete plucking of all teeth on the experimental side was not feasible (illustrated in Figure 1A); accordingly, only sufficiently erupted teeth were removed, while smaller or partially erupted teeth remained.

**Figure 2.**
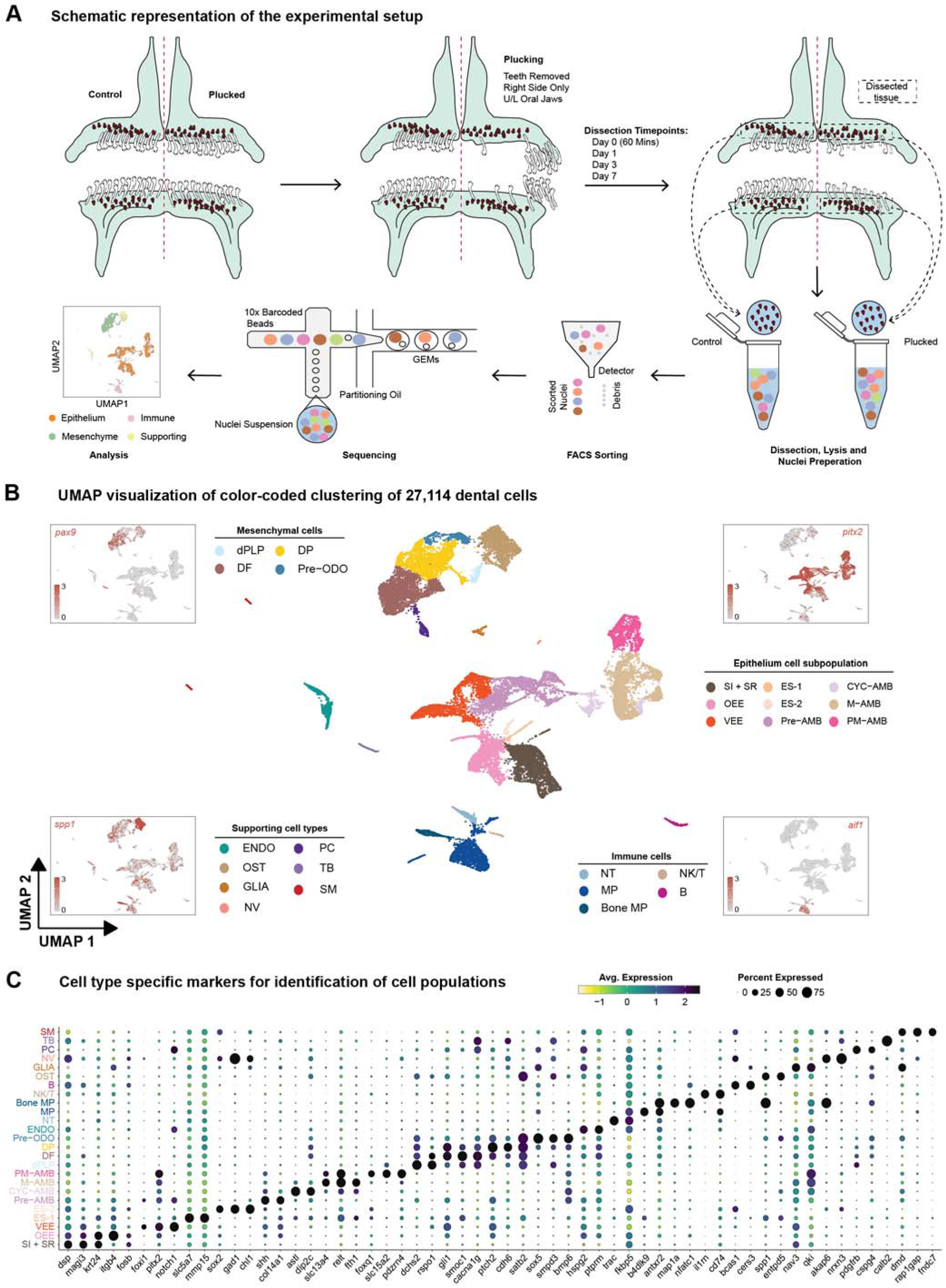
Single-nucleus RNA-seq of cichlid replacement teeth. A. Schematic of the snRNA-seq experimental and processing workflow. B. Uniform manifold approximation and projection (UMAP) of 27,114 nuclei. Points that are close together represent nuclei with similar gene expression profiles. The color of each point represents the cell type. SI + SR, Stellate reticulum and Stratum intermedium; OEE, Outer Enamel Epithelium; VEE, Ventral Enamel Epithelium; ES-1, Epithelial Supporting Cells 1; ES-2, Epithelial Supporting Cells 2; Pre-AMB, Pre-Ameloblasts; CYC-AMB, Cycling Ameloblasts; M-AMB, Maturing Ameloblasts; PM-AMB, Post Maturation Ameloblasts; dPLP, Distal Pulp; DF, Dental Follicle; DP, Dental Papilla; Pre-ODO, Pre-Odontoblasts; ENDO, Endothelium; NT, Neutrophils; NV, Nerves; MP, Macrophages; Bone-MP, Bone Macrophages; NK/T, Natural Killer and T Lymphocytes; B, B Lymphocytes; OST, Bone; GLIA, Glia; PC, Pericytes; TB, Taste Buds; SM, Smooth Muscle Cells. Canonical marker genes used to identify major cell classes include *pitx2* (dental epithelium,) *spp1* (bone), *pax9* (dental mesenchyme), and *aif1* (immune) (Supplementary Table 1). C. Expression of marker genes used for identification of major cell types in (B)

Manipulated individuals of CA, MZ Red and PT differed in the number of new teeth on the control and plucked sides of the jaw, consistent with previous observations of divergent total tooth numbers in these species^30^ (Supplementary Figure 3). To account for differences between species, we calculated ‘tooth gain’ as the number of new teeth divided by total teeth on each jaw quadrant in each individual. Notably, tooth gain was ∼3-4x higher on the plucked side of the jaw, across all three species (Figure 1C-D; Supplemental Figure 3). Approximately 60% of teeth were new, on the plucked side, across species, after 15 days. We conclude that plucking accelerates tooth replacement on the plucked side, with similar tempo and mode across Malawi cichlid species with divergent dentitions.

### snRNA sequencing reveals conserved dental and non-dental cell types

To identify cell types and molecular signals involved during early stages of accelerated tooth replacement, we collected snRNA-seq profiles, in plucked vs control jaw quadrants, at four time points. We conducted these experiments in MZ Red because (i) the cichlid genome reference comes from a closely related species in this genus, (ii) MZ Red jaws are relatively large and packed with replacement tooth germs and (iii) the response to plucking was consistent across species (Figure 1C-D). Upper and lower jaws of each study animal were split into control and plucked sides, with teeth plucked on the experimental (right) side under anesthesia. After teeth were plucked, fish were returned to isolated water tanks to recover, after which oral jaws were collected at four different times: day 0, i.e. 60 minutes; 1 day; 3 days; and 7 days after plucking (Figure 2A). As above, individuals act as their own control in these experiments; we conducted snRNA-seq on 2-4 replicates per time point. Replicate samples were pooled by condition at each time point and later deconvoluted using genetic variants in snRNA-seq reads.

In contrast to our previous approach of dissecting individual replacement teeth^24^, here we collected the minimal jaw region harboring replacement teeth along with surrounding tissue, with the intent to sample not only dental but also supporting cells that may contribute to the replacement process (Figure 2A). In total, ∼3 billion RNA-seq reads were sequenced and aligned to the Lake Malawi cichlid *Maylandia (Metriaclima) zebra* reference genome^34^. After data pre-processing, 27,114 nuclei passed quality control filters. These high-quality nuclei were then deconvoluted using souporcell^35^, which clusters cells using genetic variants detected within the snRNA-seq reads and assigns them to inividuals. Data quality was consistently strong across individuals, time points and conditions and thus supported consolidation into a single UMAP for cluster delineation and cell type identification (Figure 2B; Supplementary Figure 4).

Following data normalization, dimensionality reduction, and clustering, cell type identities were inferred using both unsupervised and supervised approaches. In the unsupervised analysis, SAMap^36^, a computational tool designed for comparing cell types from distantly related species while accounting for protein sequence divergence, was used to integrate our snRNA-seq data with existing scRNA-seq data sets from mouse incisor^21,22^ and identify homology across cell types with shared gene expression patterns (Supplemental Figure 5). This was followed by a supervised analysis using the expression of canonical marker genes from the literature to further validate cell types^20–22,24^; this includes our own previous work with *in situ* hybridization and immunohistochemistry in cichlid teeth^1,10,11,30,31^. Nuclei were annotated at both coarse- and fine-grained scales, facilitating the identification of 25 primary cell types (n=25) containing ∼60-2,900 nuclei and 50 secondary clusters (Supplemental Figure 4), subsequently classified into four cell classes (n=4) containing ∼100-13,000 nuclei (Figure 2B; e.g., epithelial, mesenchymal, immune, supporting).

The dental epithelial compartment, marked by expression of *pitx2*, was further subdivided into canonical cell types: outer enamel epithelium (OEE) characterized by the expression of *krt15, fosb, itgb4, and egr1*; stratum intermedium and stellate reticulum (SI + SR), marked by the expression of *dsp, magi3,* and *cgnl1*^20^; the ventral or labial extension of the outer enamel epithelium (VEE)^10,11,22^, marked by expression of *notch1* and *foxi3*; and the lineage of ameloblasts marked by expression of *odam*. We delineated four ameloblast cell types corresponding to different stages of differentiation: pre-ameloblasts (pre-AMB); cycling ameloblasts (CYC-AMB); maturing ameloblasts (M-AMB); and post-maturation (PM-AMB) ameloblasts (Figure 2C; Supplemental Figure 5).

We identified two transcriptionally distinct but numerically small cell populations, adjacent to OEE and SI + SR clusters, which we named epithelial supporting cells, ES-1 and ES-2, respectively. ES-1 cells are enriched for expression of *slc7a5* and *mmp15* (Figure 2C); this cell type does not have a best match in comparison to mouse incisor epithelium or mouse incisor cell types (Supplementary Figure 5). ES-2 cells are enriched for *sox2*, *gad1*, and *chl1*, all of which are expressed in oral and dental epithelium in cichlid tooth and taste bud development^11^. ES-2 cells are most similar to mouse incisor OEE and SI + SR (Supplementary Figure 5).

The dental mesenchyme, marked by *pax9* and *pdgfra* was further subdivided into: dental follicle (DF) characterized by *smoc1*, *gli1* and *cacna1g*; dental papilla (DP), with increased expression of *ptch2*, *cdh6*, *egr3*, and *fgf3*; and pre-odontoblasts (pre-ODO) enriched for expression of *satb2, sox5, smpd3*, and *bmp6*. We also identified a cluster annotated as distal pulp (dPLP), marked by *dchs2*, *rspo1* and *lgr5*.

We annotated a number of other cell types, sometimes in low numbers, found to be associated with mammalian teeth, including glia (GLIA), alveolar bone (OST), pericytes (PC), immune cells (NK/T, macrophages [MP], etc.) and endothelium (ENDO)^20–23,37^. Finally, due to our method of collecting jaw tissue around replacement teeth, we also identified cell types characterized by tastebud (TB), nerve (NV), and smooth muscle (SM) markers. Overall, these data support the conclusion of deep conservation of cell types and cell type gene expression across vertebrate teeth^24^.

### Epithelial developmental trajectories bifurcate from the presumptive successional lamina

One of the remarkable features of snRNA-seq experiments from complex tissues, realized by Trapnell and colleague more than a decade ago^38^, is that a single-timed experiment (or the combination of experments over timepoints) contains cells or nuclei across a continuum of differentiation states. When that continuum is inferred, cells are ordered along ‘pseudotime’ trajectories and features of differentiation can be estimated. We sought to determine developmental trajectories in the epithelial cell types of cichlid replacement teeth for two reasons. First, this has been done in mouse incisors^21–23^ and we wished to know if trajectories are similar. Second, we aimed to identify clusters of cells expressing markers of the cichlid successional lamina^10,11,31^, the band of epithelium that initiates tooth replacement. To gain power in this analysis, we pooled cells across time points and experimental conditions after demonstrating that each epithelial sub-population was observed in each condition at each time point, and that inferred trajectories did not differ in plucked vs control conditions (Supplemental Figure 6).

To infer developmental potential of epithelium-derived cells, we employed CellRank 2 with a CytoTRACEKernel^39^. This method calculates transition probabilities for nuclei based on the connectivity of the k-nearest neighbor graph and CytoTRACE scores; the approach determines the direction of cellular state changes, which is beneficial when the initial state is unknown. Our analysis identified a consistent flow from the VEE, branching into two distinct lineages: (1) consisting of ameloblasts at various stages of differentiation, and (2) involving an epithelial subgroup including OEE, SI + SR, and ES-1 and ES-2 cells (Figure 3A).

**Figure 3.**
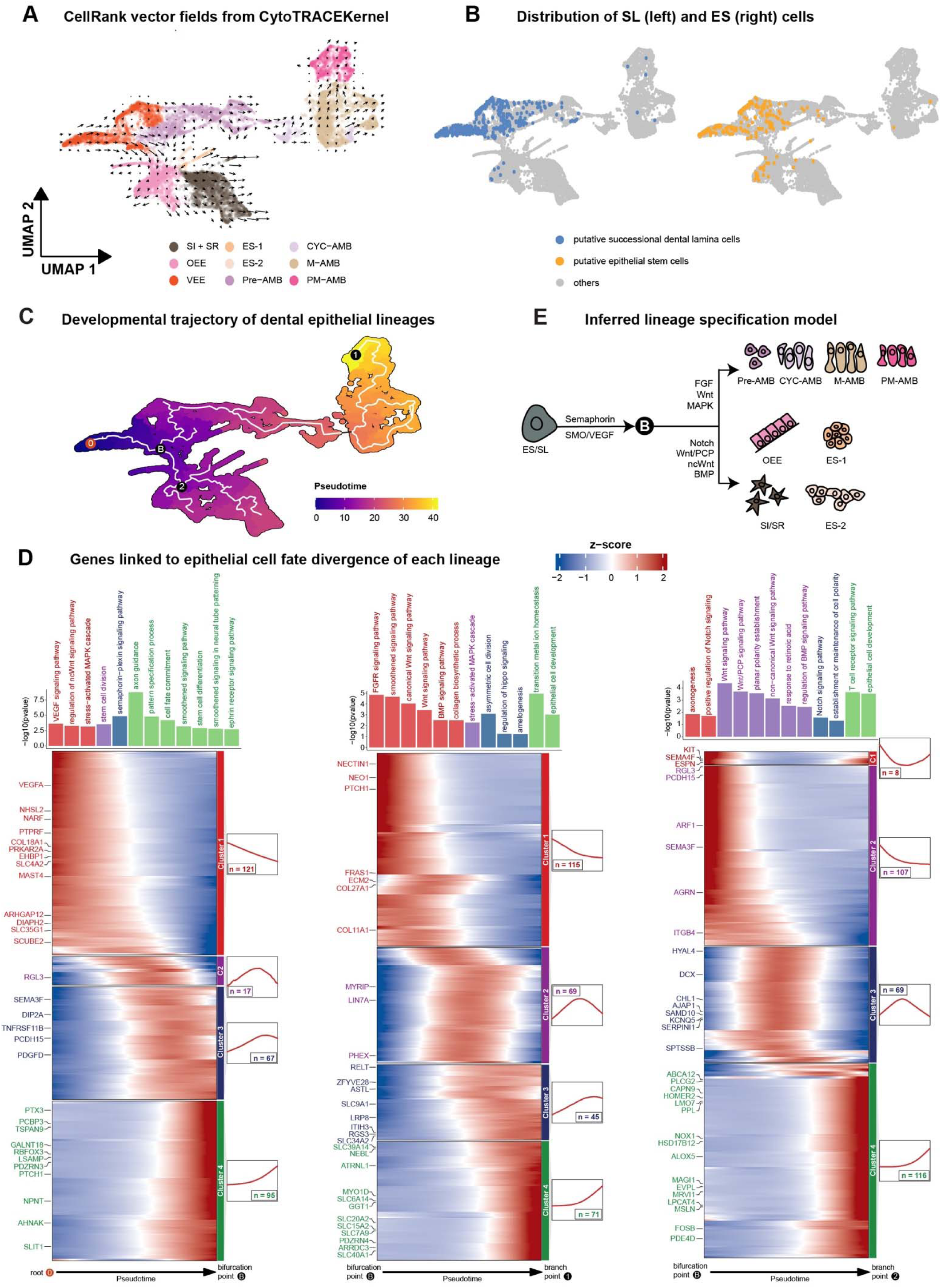
Transcriptional programs of developmental potential in replacement tooth epithelium. A. CellRank 2 projected velocity fields showing inferred cell–cell transition dynamics in epithelium using CytoTRACE-derived developmental potential. Arrows indicate the predicted direction of state change in the UMAP embedding. B. UMAP overlay with SL (left) and ES (right) cell distribution, identified by co-expression of canonical marker genes curated from the literature. C. The developmental trajectory of dental epithelial cells is analyzed via pseudotime alignment. D. Gene expression dynamics along pseudo-differentiation reveal distinct transcriptional modules, with corresponding GO enrichment for each module. E. Schematic diagram summarizing lineage specification from ES/SL cells.

To identify putative SL nuclei, we found cells enriched for canonical SL *(pitx2, sox2, shh, igfbp5, foxi3, bmp4, gli1)* and epithelial stem cell (ES) markers *(lrig1, pknox2, spock1, pcp4, sox2, bmi1, gli1, lgr5, igfbp5, zfp273)*^11,21,40,41^. Nuclei enriched for markers of SL and ES were concentrated in the VEE cluster (Figure 3B); this pattern notably aligned with the origin point of developmental potential inferred from CellRank 2.

To map developmental trajectories in epithelial cells of cichlid replacement teeth, we carried out a pseudotime analysis using Monocle3^42^. We set the starting point (0) in the VEE, consistent with our analyses of developmental potential and enrichment of SL/ES markers (Figure 3A-B). We observed a similar bifurcation (marked B in Figure 3C) in this analysis, with one path (B>1) leading along a differentiation trajectory of ameloblasts and a second path (B>2) leading to all other epithelial sub-clusters. The particular sequence of ameloblast differentiation we observed is as expected from studies in the mouse incisor^21,22^. We next identified co-expressed genes and their functional enrichment along each of the three main psuedotime trajectories, from 0>B, from B>1 and from B>2 (Figure 3D). Pathways enriched along the trajectory from the VEE to the bifurcation point (0>B) include VEGF signaling, MAPK signaling, stem cell division, Semaphorin-plexin signaling, Smoothened signaling and axon guidance. The ameloblast maturation trajectory (B>1) is enriched for FGF, WNT and BMP signaling as well as ‘amelogenesis.’ Finally, the trajectory from the bifurcation point to all other epithelial lineages up-regulates Notch signaling, axogenesis and planar polarity, among others (summarized in Figure 3E). Taken together, we identified cells within the epithelial compartment of cichlid replacement teeth that express markers of the successional lamina, and we demonstrated that all epithelial lineages, including an ameloblast maturation trajectory, are derived from these progenitor cells.

### Multiple developmental trajectories define the mesenchyme of cichlid replacement teeth

Work in the mouse suggests that dental mesenchyme is heterogeneous, with multiple niches containing stem cells^18,19,21^. We used a similar approach as above to identify progenitor populations and then trace developmental trajectories from those points. Once again, we pooled cells across time and experimental conditions after demonstrating that each mesenchymal sub-population was observed in each condition at each time point, and that inferred trajectories did not differ in plucked vs control conditions (Supplemental Figure 7).

Using CellRank 2, we found multiple centers of developmental potential (Figure 4A). These domains coincided alternatively with the expression of *twist1* and *twist2* genes, known to influence the proliferation and differentiation of dental MSCs^43,44^ (Figure 4B). Notably, *twist1* expression overlapped with *dnmt1*, a regulator of DNA methylation that retains the stem state of dental mesenchyme^45^, as well as *runx2*, which is enriched in dental ectomesenchyme (DEM) progenitor niche^46^. The co-expression of these factors, along with that of dental stromal markers, delineates a putative DEM progenitor domain (Supplemental Figure 8A-C). We used Monocle3 to verify and and further eluclidate these multiple trajectories, with starting nodes at the orgins of developmental potential, in the DEM (node 0), DF (node 1) and DP (node 2, Figure 4C). Multiple progenitor domains for dental mesenchyme, across DEM, DF and DP, are observed similarly from trajectory analysis of scRNA-seq in mouse and human teeth^47^. The trajectory from node 1 bifurcates into follicle (F) and pericyte (P) lineages, consistent with the known association between mesenchymal stem cells and perivascular niches^48^.

**Figure 4.**
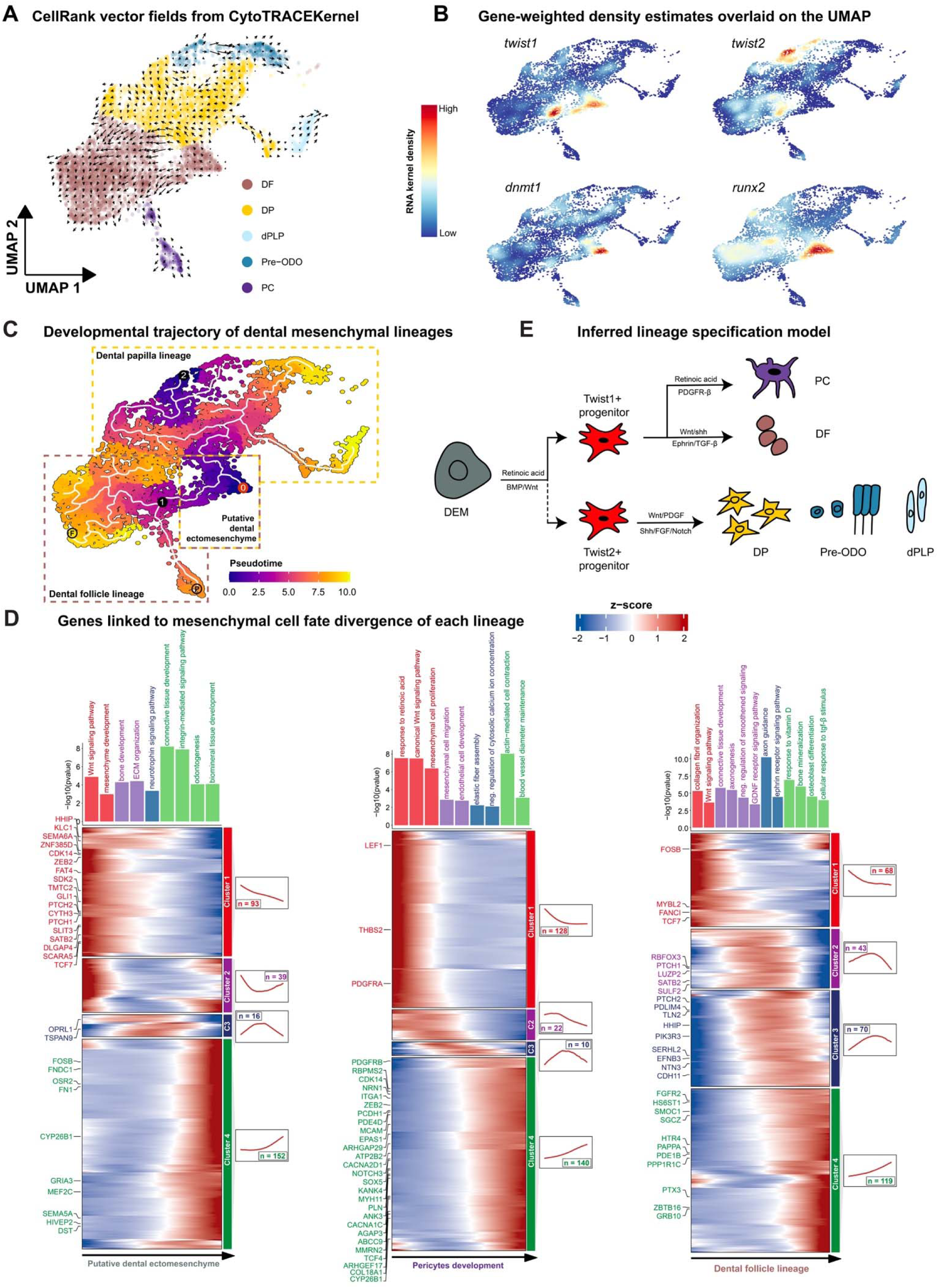
Transcriptional programs of developmental potential in replacement tooth mesenchyme. A. CellRank 2 projected velocity fields showing cell–cell transition dynamics in mesenchyme using CytoTRACE-derived developmental potential. Arrows indicate the predicted direction of state change in the UMAP embedding. B. Gene-weighted density estimates overlaid on the UMAP showing expression distributions of *twist1*, *twist2*, *dnmt1*, and *runx2* across mesenchymal states. C. The developmental trajectory of mesenchymal cells is analyzed via pseudotime alignment. D. Gene expression dynamics along pseudo-differentiation identify distinct transcriptional modules, with corresponding GO enrichment for each module. E. Schematic diagram summarizing lineage specification from mesenchymal progenitor cells.

Across the DEM developmental trajectory, we observed early expression of markers involved in BMP and WNT signaling, followed by divergence into odontogenic programs corresponding to DF and DP lineages, respectively (Figure 4D). Differentiation from the origin of the developmental trajectory in the DP involved gene expression signatures of FGF, Smoothened, WNT, Semaphorin and PDGF signaling as well as ‘odontogenesis’ (Supplemental Figure 7D); whereas the DF trajectory was characterized by WNT, TGF-/3, and Ephrin signals (Figure 4D). Along the pericyte lineage, we observed enrichment of PDGFR-/3 and RA signaling (Figure 4D). Overall, we conclude that multiple developmental trajectories act to specify cell types in the mesenchymal compartment. Each of these originates in a region of developmental potential marked by a distinct *twist* gene, *dnmt1*, and *runx2*, followed by different patterns of combinatorial signaling along distinct differentiation paths (Figure 4E).

### Plucking induces a distinct time course of gene expression during accelerated tooth replacement

To identify signatures of accelerated tooth replacement, we collected snRNA-seq profiles, in replicate plucked vs control jaw quadrants, at four time points. After teeth were plucked, fish recovered in isolated water tanks, and oral jaws were collected at four times: day 0, i.e. 60 minutes; 1 day; 3 days; and 7 days after plucking (Figure 2A). Individuals act as their own control in these experiments. We assessed four features, quantifiable with single cell data, that may differ on experimental (plucked) vs. control sides of the jaw. Plucking might change the: (1) composition of cell types; (2) ‘state’ of cells in cell types; (3) expression of genes in cell types and/or (4) interactions amongst cell types.

We used two approaches to to infer compositional differences amongst cell types on plucked vs. control jaw halves. scCODA^49^ and Cacoa^50^ make different assumptions and use different models to ascertain compositional shifts. We observed modest, consistent (across approaches) compositional differences between plucked and control conditions, confined to epithelial cell types. For instance, the cell type SI + SR was observed in greater proportion in plucked samples, on day 0 and day 1. The cell type VEE was observed in greater proportion in plucked samples on day 3 while SI + SR and OEE were in greater proportion in control samples, as that time point (Supplemental Figure 9). We used CytoTRACE^51^ to test whether cell types differed in cell state (e.g., developmental potential) across conditions and found no difference in cell state between plucked vs. control conditions (Supplemental Figure 10). We surmise that lack of dramatic differences in cell type compostion and cell state reflect our manipulation, which is not an ablation^22^, but rather an acceleration of a common process on the plucked side.

We identified considerable variance in cell-type specific gene expression that itself varies with time (Figure 5A). Thus, we modeled joint effects of time and condition and observed differentially expressed genes (DEGs) across the time course of tooth replacement after plucking (Figure 5B). Overall, the pattern is of DEG up-regulation on the plucked side one hour after manipulation, especially in epithelial cell types, with more balanced up- or down-regulation on the plucked side, to follow on days 1, 3 and 7 (Supplementary Table 2).

**Figure 5.**
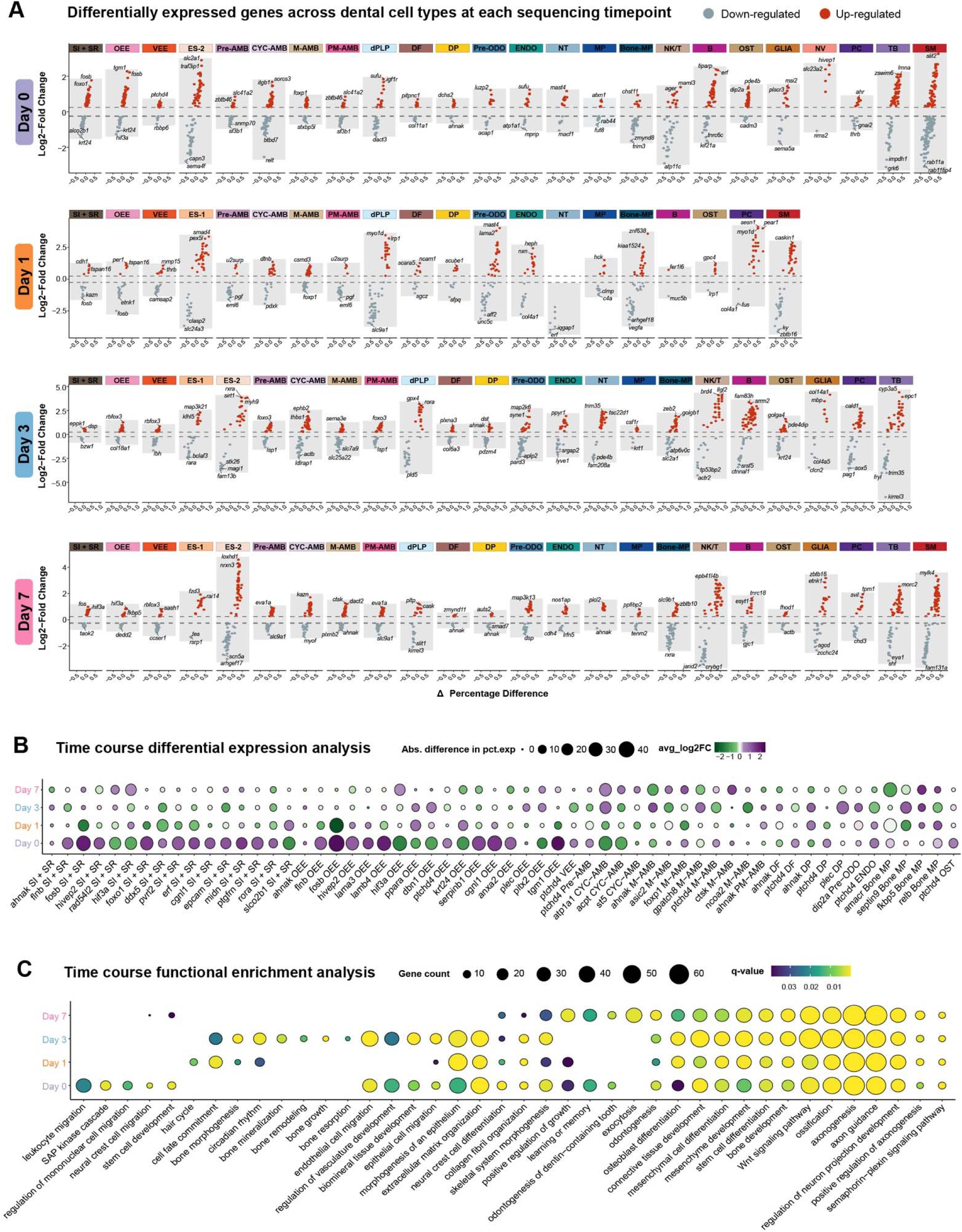
Gene expression dynamics across cell types following tooth plucking. A. Differentially expressed genes identified within each cell types, comparing plucked to control across time points. Red and blue dots highlight the top 2-3 up-regulated and down-regulated DEGs, respectively. B. Stage-resolved expression dynamics of DEGs across the time course. The size of each dot is proportional to the percentage of cells expressing DEGs. Purple and green dots indicate DEGs showing higher and lower expression, respectively, in plucked, as compared to control. C. Time-course GO biological processes reveal enriched functional profiling at each developmental stage. The size of each dot is proportional to the number of genes contributing to each enriched term, and the color scale indicates statistical significance.

A few genes with dynamic expression patterns are worth mentioning. *fosb* is strongly upregulated one hour after manipulation (day 0), on the plucked side, in OEE and SI + SR cell types, but then down-regulated in these same cell type compared to control at day 1. *Fos* was found to be a marker of OEE progenitors in mouse incisors^21^. *foxo1*; a transcription factor essential for enamel maturation^52^ exhibits a similar expression profile in SI + SR. Notably, a set of genes including *pitx2*, one of the earliest markers of tooth development, is down-regulated in plucked samples at day 0 within the OEE, but up-regulated at later timepoints. Similarly, *atp1a1* and *acpt*, both involved in late-stage enamel formation^53,54^, are down-regulated at early timepoints but elevated by day 7, in cycling ameloblasts of plucked samples. Finally, *ptchd4*, a mediator of Hedgehog signaling, exhibited a near-identical expression profile across multiple cell types; first upregulated on the plucked side at day 0, down-regulated in plucked at day 3 and then up-regulated again at day 7.

Functional enrichment analysis of DEGs by day similarly identified processes unique to specific time points and/or processes common to most. For example, day 0 DEGs were enriched for aspects of inflammation, immune response, neural crest and stem cell biology, while most time point DEGs were enriched for epithelial and mesenchymal development, odontogenesis and axon guidance (Figure 5C).

### Dynamic cell-cell communication accompanies accelerated tooth replacement

Tissue interactions are central to tooth development^55,56^. Lumsden’s classic tissue recombination experiments identified reciprocal crosstalk between oral epithelium and neural crest derived mesenchyme, as well as changing sources of odontogenic potential during tooth development. Because these interactions are so well characterized, and because of the dynamic patterns of celltype-specific gene expression during accelerated tooth replacement (Figure 5), we asked whether we could detect differences in cell communication, between plucked and control conditions across time. To this end, we used CellChat^57^, which models coordinated expression of genes encoding known ligand-receptor pairs and cell adhesion binding protein pairs, to estimate the strength of interaction between cell populations. We executed CellChat for each time point post plucking, and illustrated cell type interactions in a heat map that compares plucked to control conditions (Figure 6).

**Figure 6.**
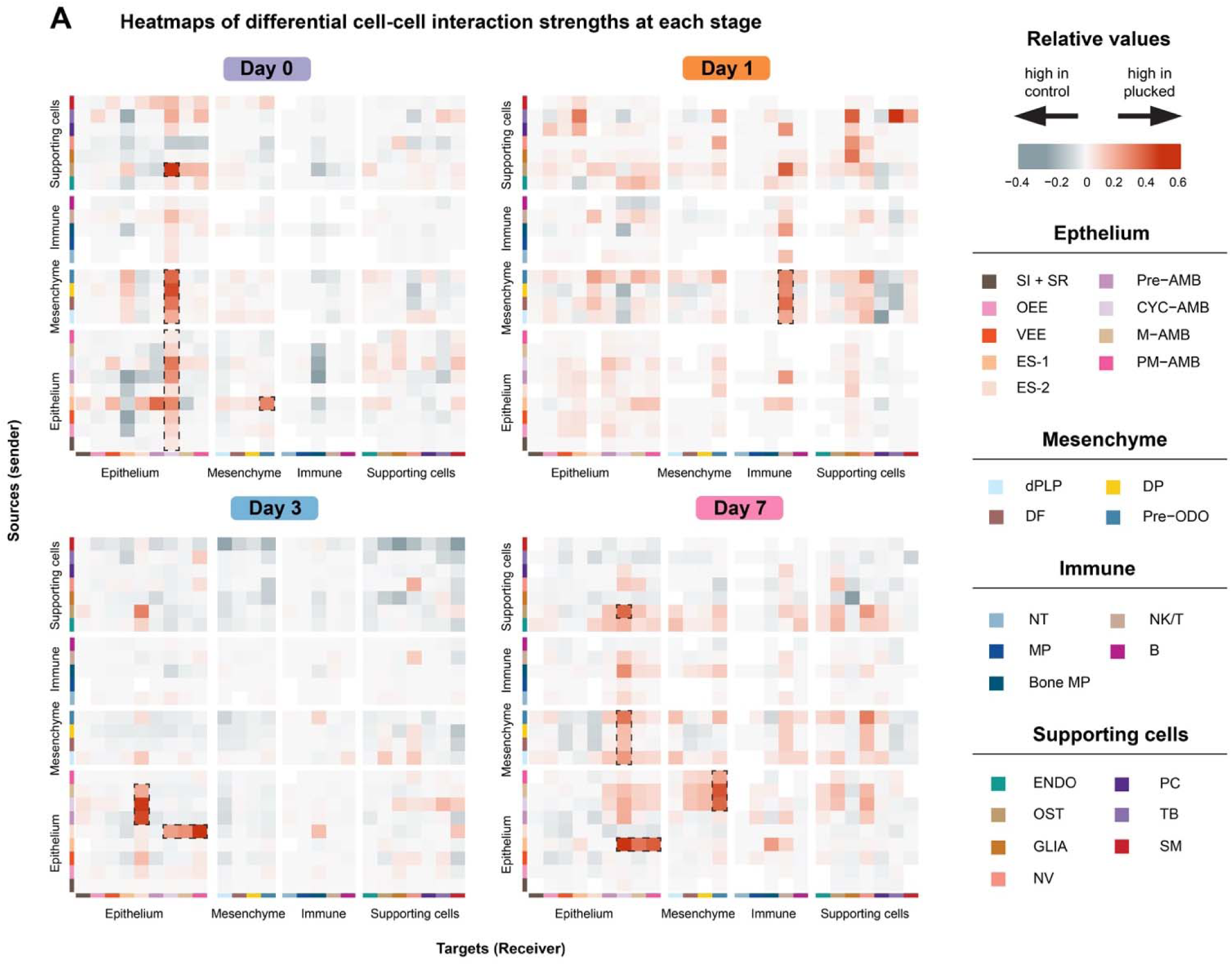
Cell-cell communication during the time course of accelerated tooth replacement. A. Heatmap of differential interaction strength among cell populations between plucked and control shows the outgoing and incoming signaling change associated with each cell group. Red (positive values) represents increased signaling in plucked and blue (negative values) represents decreased signaling in plucked, between cell groups, as compared to control. The color scale indicates the magnitude of differential interaction strength.

These data are striking in their specificity. We focus on interactions estimated to be stronger (red in the heatmap) on the plucked side of the jaw. At day 0 (60 minutes) signals were sent primarily by the mesenchymal complex (pre-ODO, DP, DF, and dPLP) and alveolar bone (OST) to CYC-AMB (see also Supplemental Figure 8D). Epithelial subpopulations pre-AMB, VEE and ES-1 also sent signals to the predominant receiver; CYC-AMB, and CYC-AMB signaled to itself. ES-1 transmitted signals to pre-ODO. On day 1, mesenchymal cell types remained the strongest source of signal, but the receiver changed from CYC-AMB to NK/T cells. Day 3 was characterized by crosstalk between epithelial cell types. Specifically, ES-2 received signals from VEE, pre-AMB, CYC-AMB, and M-AMB, while sending signals to CYC-AMB, M-AMB, and PM-AMB. On day 7, we observed (as in day 0) strong signaling from mesenchymal sub-clusters and alveolar bone (OST) to CYC-AMB. Additionally, pre-ODO received incoming signals from multiple ameloblast cell types. Day 7 signaling from ES-1 also resembles day 0, and within the epithelium seems to replace day 3 signals from ES-2. The changing and recurrent nature of cell-cell communication across time since tooth plucking likely reflects rapid morphogenesis of pioneer replacement teeth as well as subsequent development of next-in-line tooth buds.

After identifying cell populations that putatively interact during accelerated tooth replacement, we wanted to explore signaling pathways that orchestrate this crosstalk. We therefore focused on enrichment of signal in celltype interaction pairs (Figure 6), highlighting pathways biased towards control or plucked experimental conditions (Figure 7A). The overwhelming signature in these data is enrichment of growth factors, components of the extracellular matrix (ECM) and pathways involved in immunity, angiogenesis, axon guidance and nerve growth. Regardless of time since plucking, cell-cell communication on the plucked side is differentially driven by Collagen, MMP (matrix metalloproteinase), Slit-Robo, SPP1 (Osteopontin), Semaphorin, and Notch signals (see also Supplemental Figure 8E).

**Figure 7.**
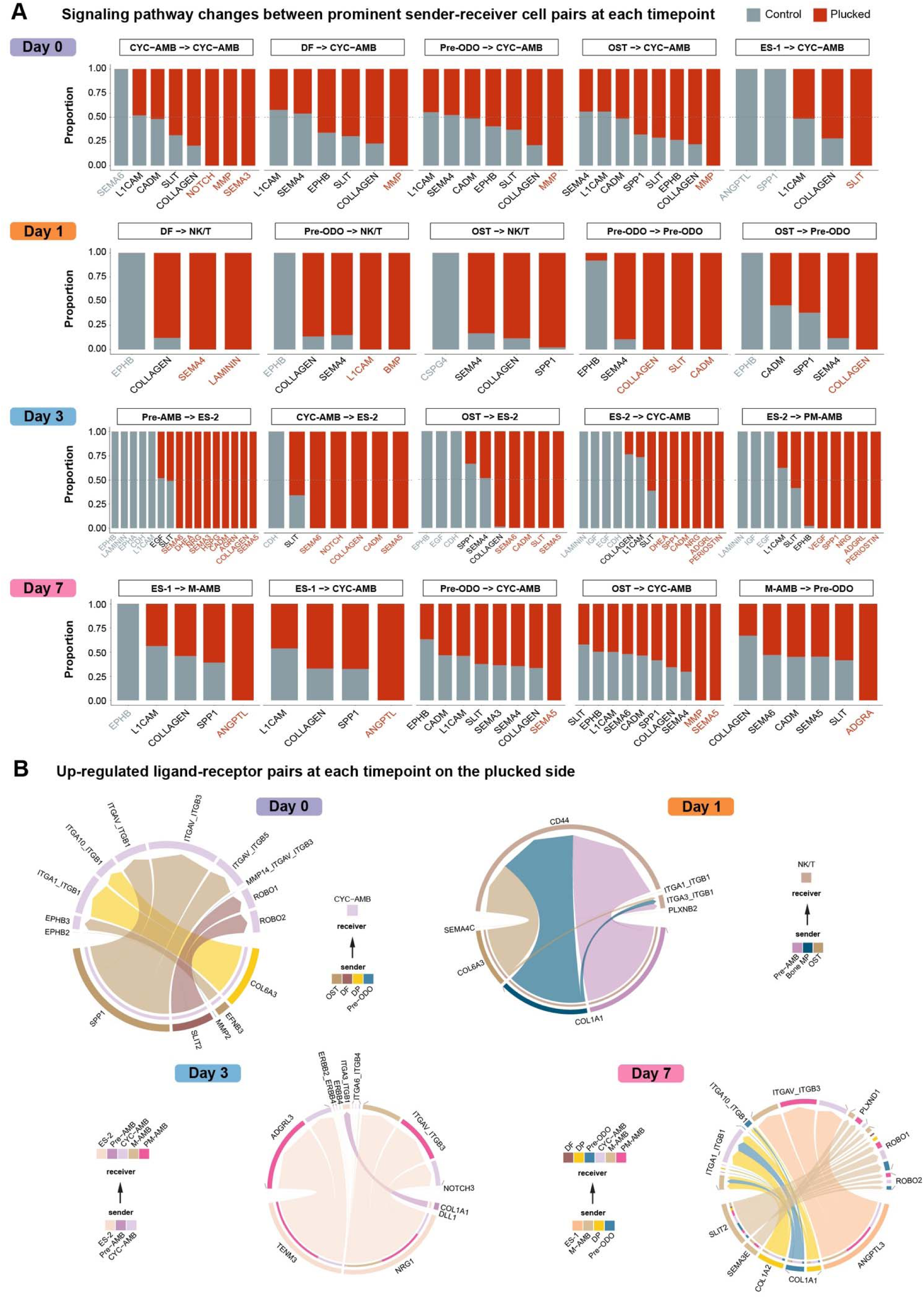
Signaling pathways and ligand-receptor interactions during accelerated tooth replacement. A. Signaling pathways between each sender-receiver cell type pair were ranked by differences in communication probability between conditions. Pathways detected exclusively in plucked are shown in red, whereas pathways detected exclusively in control are shown in blue. Pathways shown in black were detected in both conditions but exhibited significant differences in communication probability between conditions. B. Up-regulated ligand-receptor interaction network in plucked compared to control. Signaling sources (senders) and targets (receivers) are shown on the bottom and top, respectively. Colored segments indicate their cell identity. Segment size is proportional to the total outgoing or incoming interaction strength associated with each ligand-receptor pair in the corresponding cell sub-population.

Notably, the ligand-receptor pairs that underlie enriched pathways are both consistent and divergent across the time course of accelerated replacement (Figure 7B). For example, at day 0 when mesenchymal cell types signal to CYC-AMB, collagen ligands pair with integrin receptors, indicative of function in the ECM. However, when these senders interact with NK/T cells at day 1, collagen ligands signal to CD44, indicative of immune response. These factors are important in tooth morphogenesis^58–60^ and reflect the scope of (i) tissue remodeling (bone and tooth), (ii) tooth growth/development and (iii) re-wiring of tooth germs to the nervous system, across the plucked jaw-half.

## DISCUSSION

Teeth are ancient vertebrate structures^61,62^ that exhibit dramatic variation in size, shape, number, location and attachment mode, across lineages. In their long evolutionary history, teeth, like other ectodermal organs (hair, feathers, scales) retained an ancestral capacity for lifelong replacement. And yet, in part because mice do not replace teeth at all, we lack a full understanding of the process. We sought to interrogate the tempo and mode of tooth replacement across dental-diverse Lake Malawi cichlid fish species^30,63^, using a plucking paradigm and a pulse-chase method to track new teeth. We demonstrated that plucking accelerates tooth replacement, only on the plucked side of the jaw. We then determined celltype-specific gene expression profiles and interactions amongst celltypes that accompany accelerated whole-tooth replacement.

### Plucking-accelerated tooth replacement has a limit

Plucking of functional teeth on the right jaw quadrant accelerates tooth replacement ∼3x, only on the plucked side. This is superficially similar to (i) incisor clipping in mice, which is followed by accelerated growth only on the clipped side^18^, and (ii) intermediate-density plucking of hairs, which initiates a synchronized, quorum-level response to organ regrowth^25^. It is different than removal of second-generation teeth in geckos, which slows the rate of replacement^64^. These comparables are reminders that the nature of the manipulation matters (e.g., functional teeth vs. second-generation teeth) as does the scope or degree of manipulation (low-density hair plucking does not initiate a quorum response^25^). This accelerated rate of ∼3x was strikingly consistent across closely related species with widely divergent dentitions (tens of large unicuspid teeth vs. hundreds of small multicuspid teeth) suggesting an invariance or constraint, which may be developmental and/or physiological.

### Identification of epithelial and mesenchymal developmental trajectories in replacing dentitions

We aimed to understand the repertoire of cell types and their developmental trajectories in cichlid replacement teeth, comparing these to mammalian counterparts that are divergent in evolutionary time (hundreds of millions of years) and capacity for regeneration. The main result of strong cell type conservation, in both epithelium and mesenchyme, as well as accessory cell types, supports earlier work^24^.

In the epithelium, we inferred a single population of cells likely to act as progenitors. This cell population was enriched for markers of the successional lamina and was most closely related to the mouse incisor VEE (ventral part of enamel epithelium). Developmental trajectories from ‘VEE’ bifurcated to give rise to (i) ameloblast cell types differing in maturation state and (ii) all other epithelial lineages, including OEE and SI + SR. This trajectory of ameloblast maturation is similar to that observed in mouse incisors^22^ and human teeth^23^, but the overall relationship amongst epithelial cell types is different from mammals, which in turn differ from one another. Future experiments should target common methods and developmental stages, in additional species (sharks, reptiles), to better understand the evolution of dental epithelial cell types.

In the mesenchyme, we observed multiple progenitor populations, one in putative dental ectomesenchyme (DEM) and then one in the dental papilla (DP) and dental follicle (DF). Each of these progenitor populations is marked by a distinct *twist* gene, *runx2* and *dnmt1*, respectively. This pattern is similar to dental mesenchyme in the human molar, where DEM bifurcates to form DF and DP lineages which further undergo fate specification, with the DP being the major source of the odontoblast lineage^23^. Multiple progenitor domains for dental mesenchyme are also observed in mice^47^.

The evolution of vertebrate dental cell types and their hierarchical relationships is a new and notably incomplete endeavour. And yet, comparison of mammal to fish dental cell types appears to follow a central tenet of evolutionary developmental biology – strong evolutionary conservation of building blocks, even when the ultimate structure or function is distinct. We showed some time ago that all teeth in the history of dentitions, regardless of location on oral or pharyngeal jaws, likely share a core gene network for tooth initiation^1^. It is tempting to infer a similarly conserved, core toolkit of vertebrate dental cell types.

### Time-sequenced cellular profiling of tooth regeneration

Regenerating organ systems share at least four features: (i) activation, (ii) initiation of regenerative molecular programs, (iii) coordination with supporting cell types, and (iv) control of size through morphogenesis^65^. We carried out time-sequenced cellular profiling of accelerated tooth replacement and captured signatures of these ‘regeneration rules’ in patterns of gene expression and cell communication (Figures 5-7, summary in Figure 8).

**Figure 8.**
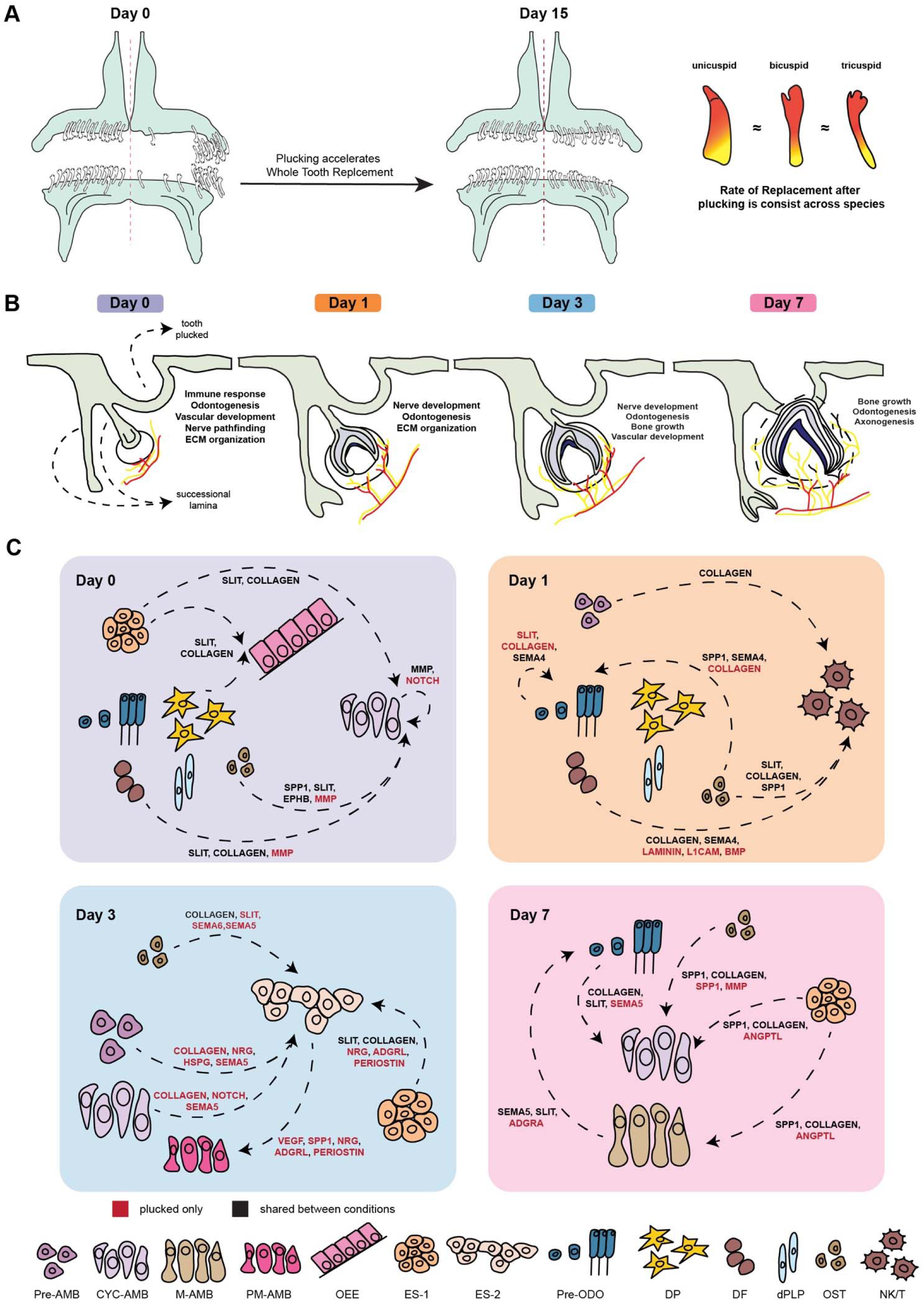
Summary of accelerated tooth replacement following plucking. A. Plucking accelerates tooth replacement, only on the plucked side, with a ∼3-4x increase in replacement rate that is conserved across cichlid species with widely divergent dentitions. B. Conceptual timeline of biological processes activated following tooth removal that accompany accelerated tooth replacement. Early immune activation is associated with increased axonogenesis and vasculogenesis, followed by elevated odontogenesis and bone formation. C. Schematic representation of prominent intercellular communication events and signaling pathways associated with accelerated tooth replacement after plucking.

Plucking activates dynamic celltype-specific gene expression, almost immediately. Genes are differentially expressed in nearly all cell types at day 0 (60 minutes after plucking) and up-regulated DEGs are enriched for function in inflammation and immune pathways. Notably, one of the strongest up-regulated genes at day 0 was *fosb*, in cell types OEE and SI + SR. *Fos* was identified as a novel marker of OEE progenitors in mouse incisors^21^ and has long been known as an immediate-early gene (a marker of plasticity) in the nervous system^66^. Fos genes may be general sentinels of cellular activation and plasticity.

Day 0 is characterized by concerted signaling from mesenchymal cell types and alveolar bone to CYC-AMB on the plucked side, likely reflecting the mesenchyme’s known role directing (replacement) tooth morphogenesis^55,56^. By day 1 post-plucking, these same mesenchymal cell types switch their cellular receivers from CYC-AMB to NK/T cells on the plucked side, reinforcing the role of immune function in tooth replacement. The immune system is not well studied in tooth development, but this is a central component of hair follicle regeneration^25,67^. By day 7 post-plucking, we observe restoration of mesenchymal signaling to CYC-AMB, and epithelial signaling to pre-ODO, on the plucked side. The latter may reflect epithelial control of next-in-line tooth bud initiation, as the primary replacement tooth continues morphogenesis and eruption.

Throughout the time course of accelerated tooth replacement, cell-cell communication on the plucked side is dominated by Collagen, MMP (matrix metalloproteinase), Slit-Robo, SPP1 (Osteopontin), Semaphorin, and Notch signaling, clear indicators of tissue remodeling, immune response, tooth morphogenesis and growth, angiogenesis and nerve pathfinding (Figure 8).

### Regeneration as ‘reinventing’ ancestral developmental programs

Recent work has explored the degree to which ancestral regeneration programs lost in humans stem from cell type extinction or modification of cell type function. In bower-building cichlid fishes, brain cell type proportions are modulated by adult neurogenesis, mediated by radial glia, cells that serve a similar function in mammalian embryos^68^. Rabbits are capable of ear regeneration and mice are not because of retinoic acid signaling differences in conserved wound-induced fibroblasts^69^. Similarly, thymus regeneration in axolotls is driven by midkine signaling in conserved thymic epithelial cells^70^. Lifelong tooth replacement follows this trend; the SL is recognizably similar to epithelial cell types in mouse and human teeth, but it no longer persists into adulthood in most mammals. Reinventing regenerative potential, therefore, might require re-engaging ancestral molecular profiles in extant cell types.

### Study limitations and future direction

This is the first manipulative characterization of whole-tooth replacement using snRNA-seq. Our work here is based on decades of histology, in situ hybridization and immunohistochemistry in the Malawi cichlid dentition^1,11,29–31^. However, these experiments would benefit from a spatial resource, providing tissue-level and temporal context to quantitative cell type gene expression and predictions of cellular communication. This is difficult in jaw tissue replete with bony crypts housing asynchronously replacing tooth families, but developments in sequence-based spatial transcriptomics should allow modification for mineralized tissue in the near future^71^. This would allow direct *in situ* interrogation of molecular and cellular profiles on plucked vs control jaw halves, over the time course of tooth replacement. Armed with information about the cell types, DEGs and signals involved in accelerated tooth replacement, future experiments should aim to manipulate these particulars of the system. We are keen to better understand the time-keeping mechanism that synchronizes tooth replacement in even- vs. odd-numbered tooth families, interactions between dental and supportive tissues (e.g., alveolar bone), and the immune environment in jaws that experience constant remodeling. Finally, we anticipate that this plucking experimental paradigm, followed by cellular characterization (snRNA-seq and snATAC-seq), can be applied to other non-model systems with lifelong whole-tooth replacement.

## METHODS

### Subjects

Cichlids were housed in the Department of Animal Resources at the Georgia Institute of Technology (Atlanta, GA) and all experiments were conducted in accordance with institutional animal care and use policies (IACUC protocol no. A100003/A100029). A total of n=25 adult cichlids were used for pulse-chase experiments, including *Metriaclima estherae* (MZ Red; n=8), *Cynotilapia afra* (CA; n=8), and *Petrotilapia* “Thick Bar” *chitimba* (PT; n=9). An additional 10 adult *Metriaclima estherae* (MZ Red) individuals were used for single-nucleus RNA sequencing experiments. Environmental conditions of aquaria were designed to approximate those of Lake Malawi. All tanks were connected to a central recirculating system and maintained at 26.7°C with a pH of 8.2. Fish were kept on a 12h:12h light:dark cycle, with full lights on from 08:00 to 18:00 Eastern Standard Time (EST) and dim lights during 60-minute transitions (07:00-08:00 and 18:00-19:00 EST). Fish were fed twice daily with Spirulina flakes (Pentair Aquatic Ecosystems, Apopka, FL, USA).

### MicroCT scanning of cichlids jaws

Prior to MicroCT scanning, rectangular foam chambers were prepared to securely hold each sample in place. Samples were removed from PBS, gently dried with a paper towel, and placed into the foam chambers. Each chamber was then inserted into a 34 mm sample holder, with the anterior of the sample faced downward into the holder and the superior of the sample oriented toward the notch of the holder. The holder was covered with parafilm, cleaned with 70% ethanol, and mounted onto the carousel of the Scanco Medical uCT50 machine. Each sample was scanned with a focus on tooth structure and the overall jaw morphology. All scans were collected using the same setting: energy/intensity at 70 kVp, 114 μA, 8 W, filter set to air, resolution set to high, FOV/diameter at 20.5, voxel size at 5 μm, integration time at 750 ms, and average data set to 1. After scanning, image data were exported as DICOM files. Segmentation was performed using the threshold feature of Materialize Magics software. Threshold values varied slightly among samples to account for differences in image contrast but were chosen to preserve tooth and jaw structures without compromising structural integrity. Edit Mask and 3D Interpolate tools were used to further refine the segmentation and to separate upper and lower jaws on the left and right sides. All microCT scans were conducted at the MicroCT Core at the Petit Institute for Bioengineering and Bioscience (IBB), Georgia Tech.

### Modified pulse-chase and tooth-plucking experiments

All individuals used in the experiments were adult fish, approximately 10-13 cm in total length with fully formed teeth and jaws. Live vital dye bone staining was performed by pulsing fish with 100 µg/ml Alizarin Red (CAS: 130-22-3, Acros Organic) buffered with 1 mM HEPES in system water for 24 hours in the dark^32^. Fish were housed individually in 3.5 L tanks on a brood rack with aeration. After the Alizarin pulse, tooth-plucking experiments were performed. For each fish, both the upper and lower oral jaws were divided along the midline into left and right halves. Teeth were removed only from the right side, which served as the experimental (“plucked”) side, while the left side served as an internal control. Plucking was restricted to an anterolateral region of the dentary, where cichlids, irrespective of their species, possess larger, evenly spaced, and morphologically distinct teeth. More posterior teeth, which are smaller (<1 mm), conical, and less phenotypically variable, were not included in the experiment (Supplemental Figure 2). Tooth removal was carried out under a stereoscope (Zeiss Stemi DV4 Stereo Microscope 8x - 32x, 000000-1018-455) using fine-toothed forceps and, when necessary, a scalpel fitted with a #11 or #12 blade. All procedures were performed under anesthesia using tricaine methanesulfonate (MS-222) in accordance with approved IACUC protocols, and care was taken to minimize tissue damage. Fish were returned to fresh anesthetic solution as needed during the procedure, with total handling time kept under 20 minutes. Following plucking, fish were returned to system water tanks for 15 days. They were then chased with 50 µg/ml Calcein (C0875-5G, Sigma) buffered with 1 mM sodium phosphate in system water for 24 hours in the dark^32^. After replacing Calcein with clean system water, the fish were allowed to recover for 24 hours before being euthanized by MS-222 overdose following previously established IACUC protocols. Upper and lower jaws were dissected from each fish, fixed overnight in 10% NBF at 4°C, and subsequently stored in 100% ethanol in the dark at 4°C. Alizarin Red labels existing mineralized bone, while Calcein labels newly mineralizing bone. Fluorescence imaging and quantification were performed using Zeiss Axio Observer Z1, and whole-mouth images were acquired using Zeiss Stereo Discovery.V12.

### Tissue Collection for snRNA Seq

All individuals used for single-nucleus RNA sequencing experiments were adult fish, approximately 10-13 cm in total length with fully formed teeth and jaws. Tooth removal was performed as described in **Modified pulse-chase and plucking**. After tooth plucking, fish were returned to isolated water tanks to recover. Oral jaws were collected at four post-plucking time points: 60 minutes (Day 0), 1 day, 3 days, and 7 days. Four biological replicates were collected for Day 0 i.e. 60-minute time point, and two biological replicates were collected for each of the remaining time points. Fish were euthanized by overdose of MS-222, after which the upper and lower oral jaws were dissected from the head. Using a stereoscope (Zeiss Stemi DV4 Stereo Microscope 8x-32x, 000000-1018-455), soft tissue labial to each jaw was carefully removed to expose the underlying jawbone. For each fish, the plucked (right) and control (left) sides of the jaw were dissected and processed separately. Replacement tooth crypts and surrounding tissue were collected by making longitudinal incision with a scalpel along the exposed jawbone, approximately 2-3 mm above and below the replacement crypt region. This approach allowed isolation of a tissue block containing all regenerating crypts with minimal surrounding tissue. Upper and lower jaws samples were rapidly dissected, flash frozen on dry ice, and stored at -80°C until preparation of single-nucleus suspensions.

### Nuclei Preparation for snRNA Seq

Nuclei were isolated using a modified Frankenstein protocol^72^ optimized for hard tissues. Immediately prior to nuclei isolation, frozen tooth-jaw samples were pooled into 2-4 biological replicates (n=2-4 replicates/pool) per condition (plucked versus control). Pools were organized by time point to balance tissue mass and nuclei yield across samples. For the 60-minute (Day 0) time point, two pools of two biological replicates each were generated and later combined during downstream analysis. For the remaining time points, pools consisted of two individual replicates with plucked and control sides derived from the same fish. Tissue masses were closely matched across pools to ensure comparable nuclei input. Frozen tissue sample pools were transferred into 500 μl of chilled Nuclei EZ lysis buffer (Sigma-Arch; NUC101). Samples were finely cut using a sterile scissor directly into lysis buffer on ice. An additional 1,000 μl of EZ lysis buffer was then added, and samples were lysed for 15 minutes with gentle nutation. Following lysis, samples were filtered using 70 μm MACS® SmartStrainers (Milltenyi) to remove large debris and remaining osseous tissue, followed by gentle trituration to achieve a homogenous suspension. Lysis was halted by adding approximately 2 ml of HBSS (Sigma-Aldrich H4385), after which samples were triturated 10-15 times using pipette tips with a 500 μm internal diameter to complete tissue dissociation. The dissociated nuclei were centrifuged (600 x g, 10 minutes, 4°C) and resuspended in approximately 1 ml of chilled wash and resuspension buffer containing 2% BSA (Sigma) and 0.2 U/μl RNase Inhibitor (Sigma) in 1X PBS (Thermo Fisher). Nuclei suspensions were filtered through 40 μm Flowmi® cell strainers (Sigma) and 30 μm MACS® SmartStrainers (Milltenyi) to remove residual debris and nuclear aggregates prior to fluorescence-activated cell sorting (FACS). Unless otherwise noted, all steps were performed at 4°C in a cold room to prevent RNA degradation.

### Fluorescence Activated Cell Sorting (FACS)

Pilot experiments indicated that passive filtration alone was insufficient to remove inorganic debris and nuclear multiplets due to the dense, mineralized nature of the tissue. Therefore, we further improved the nuclei purity and sample quality using FACS (BD FACS Melody Cell Sorter, BD Biosciences) and BD FACS Chorus Software (v1.1, BD Biosciences). Nuclei were gated using 10 μm sizing beads (BD Biosciences) and 1 μg/ml 7AAD (Sigma) to identify single nuclei based on forward scatter (size), side scatter (shape), and DNA content (7AAD fluorescence). This gating strategy effectively eliminated nuclear multiplets, irregularly shaped or damaged nuclei, and inorganic debris. Approximately 90,000 nuclei were collected per pool from both the plucked and control sides into 96-well plates (Corning Costar) containing 30 μl of wash buffer and maintained at 4°C. Sorted nuclei were used for downstream single-nucleus RNA sequencing. FACS data and gating strategies were visualized and analyzed using FlowJo v10.6.0.

### Library Preparation and Sequencing

Suspensions of isolated nuclei were loaded onto the 10x Genomics Chromium Controller (10x Genomics) at concentrations between 300-400 nuclei/μl. Target recovery was 3,000–4,000 nuclei per sample, with the goal of achieving an average sequencing depth of approximately 80,000 reads per nucleus. cDNA synthesis and library preparation were performed according to the manufacturer’s instructions using the Chromium Single Cell 3’ Reagent Kits User Guide v3.1 Chemistry (10X Genomics), including the Single Cell 3’ GEM, Library and Gel Bead Kit v3.1 and Chromium i7 Multiplex Kit. Library quality was assessed using high-sensitivity DNA analysis on the Agilent 2100 Bioanalyzer, and library concentrations were measured using Qubit 2.0 fluorometer (Invitrogen). Barcoded libraries were pooled by time point and sequenced on the NovaSeq 6000 platform (Illumina) using a single flow cell and the 300-cycle S1 Reagent kit (2 x 150 bp paired-end reads; Illumina). Sequencing was performed by the Molecular Evolution Core at Georgia Tech.

## Quantification and statistical analysis

### snRNA-seq data pre-processing and quality control

We processed FASTQ files using Cell Ranger v3.1.0. Reads were aligned to the *Maylandia zebra* (Lake Malawi cichlid) reference genome assembly^34^ using a splice-aware alignment algorithm (STAR) within Cell Ranger. Gene annotations were obtained from the same assembly (NCBI RefSeq accession: GCF_000238955.4; M_zebra_UMD2a). Because nuclear RNA contains intronic sequences, reads mapping to introns were included during the Cell Ranger count step. Unique Molecular Identifiers (UMIs) that were homopolymers, contained ambiguous bases (N), or contained any base with a quality score less than 10 were excluded. Following these steps, Cell Ranger generated filtered feature-by-barcode matrices containing expression data for a total of 32,471 features (corresponding to annotated genes) and a total of 30,005 barcodes (corresponding to droplets and putative nuclei).

Gene expression matrices generated by Cell Ranger were then assessed with Seurat v5^73^ for further data processing. To remove potentially dying, dead, or uncharacteristically stressed nuclei, barcodes with fewer than 250 total genes detected or with more than 5% of their total transcripts being mitochondrial genes were excluded from downstream analysis. To reduce the risk of doublets or multiplets, barcodes with more than 2,500 total genes were also excluded. As a result, a total of 27,114 barcodes passed all quality control filters and were included in the downstream analyses.

### Dimensionality reduction and clustering

After the log-normalization of gene expression measurements for each individual cell, confounding factors including percentage of mitochondrial genes (pct.mt) and the total number of UMI counts (nCount_RNA) were regressed out. Highly variable genes were identified using the FindVariableFeatures function in Seurat with the mean.var.plot selection method. This method aims to identify variable features while accounting for the strong correlation between variability and average expression. In this process, we pinpointed 4,000 genes that showed the most variable expression patterns across nuclei. Prior to dimensional reduction, we scaled the gene-level data using the ScaleData function in Seurat. To investigate dimensionality, we first executed a linear dimensional reduction (principal components analysis, PCA) using the RunPCA function, setting the maximum number of dimensions to 50 (dim=50). We then employed Seurat’s JackStraw, ScoreJackStraw, and JackStrawPlot functions to estimate and visualize the significance of the first 50 principal components (PCs). We also used the Elbow plot function to visualize the variance explained by these 50 PCs. Since all 50 PCs were highly statistically significant and there was no observed drop-off in variance explained across PCs, we used all 50 PCs for non-linear dimensional reduction (Uniform Manifold Approximation and Projection, UMAP) using the RunUMAP function in Seurat. For RunUMAP, minimum distance allowed apart the embedded points was set to 0.5 (min.dist=0.5), the number of neighboring points used in local approximations was set to 50 (n.neighbors=50), and Euclidean distance metric was used to measure distances between points.

Prior to clustering, we embedded nuclei into a k-nearest-neighbor (KNN) graph based on Euclidean distance in UMAP space, with edge weights based on local Jaccard similarity, using the FindNeighbors function in Seurat (k.param=50, prune.SNN=0). Finally, we performed clustering on the derived graph using Seurat’s FindClusters function, employing the Louvain algorithm with multilevel refinement (algorithm=2), to define the cluster partition. By setting the cluster resolution to 0.6, we identified 50 distinct clusters, which were then classified into 25 cell types. Cell type assignment was performed based on the combination of cluster marker gene analysis and interspecies integration of sc/snRNA-seq data from cichlids and mice.

### Cluster marker gene analysis

The biological identities of specific clusters were investigated using a combination of unbiased cluster-specific marker gene detection and a supervised examination of previously established marker genes. To find marker genes for each identified cluster, we used the Seurat function FindAllMarkers (only.pos=TRUE) with log-scaled 2-fold difference set to 0.25 (logfc.threshold=0.25, base=2) and genes detected in at least 20% of cells (min.pct=0.2) to ensure that the markers are sufficiently expressed in the corresponding cluster. Genes with Bonferroni-adjusted *p*-values < 0.05 were retained as the cluster–specific markers, based on the Wilcoxon rank-sum test. Functional enrichment analysis of Gene Ontology (GO) categories was performed on cluster-specific marker genes. Briefly, we first converted cichlid gene names to their human orthologs, and then ran enrichment analysis with ToppGene Suite under default settings. Results that survived FDR adjustment (*q*-value<0.05) were deemed statistically significant. Established dental cell-type-specific marker genes were cross-referenced with the output from FindAllMarkers to generate further insight into the biological identity of clusters.

### Interspecies integration of sc/snRNA-seq data

To further investigate and verify the biological identities of specific clusters, we integrated sc/snRNA-seq data from cichlids and mice to facilitate a comparative analysis of the transcriptional similarities among dental cell types. This analysis was conducted using SAMap^36^, which constructs a gene homology graph, weighted by sequence similarity of homologous gene pairs between species. This approach allows genes with higher sequence similarity across species to have a greater influence on the integration, making it possible to compare species with large evolutionary distances, such as cichlids and mice. First, to determine sequence similarity, we downloaded the proteomes from NCBI: *Maylandia zebra* (GCF_000238955.4) and *Mus musculus* (GCF_000001635.27). We then ran blastp to identify reciprocal blast hits between cichlids and mice using SAMap’s map_genes.sh script and constructed a gene-gene bipartite graph based on the bi-directional blastp results. Following this, we converted the raw gene expression matrices of the sc/snRNA datasets from Seurat objects to h5ad files, and processed them using the SAMAP function, specifying the gene symbols of each protein. Finally, we ran the SAMap algorithm using default parameters. Next, the k-nearest-neighbor graph produced by SAMap was used to create a similarity score between dental cell types in cichlids and mice. The similarity score was defined as the average number of k-nearest neighbors between cells from mice to cichlid nuclei. To determine significance, we performed a permutation test where we shuffled the cell type labels of cells/nuclei for both datasets 1,000 times and calculated similarity scores. Cross-species cell-type pairs with similarity scores higher than all permutations of each cichlid cell type were deemed significant. Homologous cichlid-mouse dental cell type pairs were then used to aid in the identification of corresponding cell types in our cichlid snRNA-seq data.

### Deconvolution of individuals from mixed pools of nuclei

souporcell^35^ was used to deconvolute individuals from the mixed-genotype snRNA-seq experiment pools. Briefly, souporcell clusters nuclei based on their genotype detected in their reads, where the resulting clusters reflect nuclei originating from the same individual. Before souporcell was applied, variants in each nucleus were identified from a subset of high-quality reads. First, reads were subset by those with a base quality score greater than 10 and were mapped confidently to the genome (-F 3844). Next, to deconvolute using souporcell, variants were identified using the GATK v4.1.8.1 HaplotypeCaller with default parameters, but without the MappingQualityAvailableReadFilter, to retain reads that were confidently mapped by Cell Ranger (MAPQ score of 255). Then, vartrix v1.1.22 was used for allele counting, with the parameters suggested by souporcell. To cluster nuclei by genotype, souporcell v2.0 was used with the number of clusters set to 2-4 to indicate the number of individuals/replicates in the pool and otherwise default parameters were used. These steps were repeated using a stricter threshold for the base quality score greater than 30. The output of souporcell was then compared and used to assign nuclei to individuals. Nuclei assignment was retained when both runs agreed or when either run was able to confidently assign the nucleus to an individual. Only non-doublet and assigned nuclei were left for further downstream analyses.

### Learning vector fields from CytoTRACE developmental potentials

To investigate the differentiation states of dental epithelial and dental mesenchymal cell subpopulation, CytoTRACE (Cellular (Cyto) Trajectory Reconstruction Analysis using gene Counts and Expression) v0.3.3^51^ was performed to predict developmental potential from snRNA-seq data based on the assumption that, on average, naive cells express more genes than mature cells during differentiation. The raw count matrix was supplied to CytoTRACE using the default settings. A CytoTRACE score, ranging from 0 to 1, was assigned to each cell based on its differentiation potential, with higher scores indicating greater developmental potential. Next, cell-to-cell transition probabilities were inferred using the CellRank 2 CytoTRACE kernel. Briefly, an undirected k-nearest-neighbor (KNN) graph representing cell–cell similarities was computed from the raw count matrix. The computed KNN graph, along with CytoTRACE scores, was then input into the cellrank.kernels.CytoTRACEKernel function. CellRank 2 leveraged both the local cellular neighborhood structure and CytoTRACE-derived differentiation scores to compute directed transition probabilities using the compute_transition_matrix function. Finally, the plot_projection function was used to visualize the transition matrix as vector fields.

### Trajectory inference and pseudotemporal ordering

To gain additional insights into gene expression variations during the differentiation of dental epithelial and dental mesenchymal cell subpopulations, we used Monocle3 v1.3.4^42^ to reconstruct single-cell trajectories. First, the normalized count matrix, cell type annotations, and UMAP embeddings of epithelium and mesenchyme subset in the Seurat object were imported to Monocle3 using the as.cell_data_set function. Next, trajectories were inferred with the learn_graph function using the default parameters. This step aims to fit a principal graph, using the SimplePPT algorithm^74^, on the UMAP of the data to recover underlying paths cells can take as they differentiate. Next, the order_cell function was used to calculate cell-wise pseudotime, using the root-node (beginning of the biological process) selection estimated by CytoTRACE and CellRank 2 CytoTRACE kernel. The subsequent analysis of specific branches from trajectories was performed using the choose_cells function. Within each branch, we identified genes that vary in expression over a developmental trajectory using the graph_test function, which performed the Moran’s I test on individual genes. To discover genes whose expression is significantly associated with pseudotime, we filtered the output of the graph_test function based on a threshold of *q*-value < 0.01, and then we further refined the selection by subsetting the top 300 genes with the highest Moran’s I values. Finally, pseudotime-dependent genes were clustered using k-means with ClusterGVis v0.1.1^75^, and an over-representation test was performed to test statistical enrichment for genes in each identified cluster using the genORA function in Genekitr^76^. Enrichment terms with a *q*-value of less than 0.05 were considered significant.

### Differential gene expression analysis

Differential expression (DE) testing was applied to identify variable genes across conditions. Within each cell type, differential gene expression was analyzed between plucked versus control nuclei using the FindMarkers function in Seurat with a log-scaled 2-fold difference threshold set to 0.25 (logfc.threshold=0.25, base=2). MAST^77^ hurdle model was employed as the DE test method with appropriate model covariates matching those used in the data normalization step. Genes that were significantly up-regulated or down-regulated (|log_2_FC|>0.25; *p*-value<0.05) in plucked nuclei were identified as differentially expressed genes (DEGs).

### Time-course analysis of gene expression

To further dissect these differentially expressed genes across conditions in each cell type, glmmSeq v0.5.5^78^ was applied to compare variations in gene expression over time by fitting negative binomial mixed-effects models at the individual gene level using the formula: gene expression ∼ condition*timepoint + (1|replicate). In this model, the interaction term of condition and timepoint was treated as fixed effects, accounting for variance in gene expression. In addition, this model also included a random term to account for variance explained by individual variation. To run glmmSeq with negative binomial models, we computed gene dispersion estimates using DESeq2^79^ on the raw gene count data. Following model fitting, Benjamini-Hochberg correction was then applied to the glmmseq results, with an adjusted *p*-value threshold of 0.05 to identify temporal differentially expressed genes.

### Functional enrichment analysis

To compare the functional profiles of differentially expressed genes obtained from different conditions across timepoints, a functional enrichment analysis was carried out using clusterProfiler v4.6.2^80^. Briefly, we used the compareCluster function with the biological process database in Gene Ontology (GO) to assess how enrichment results varied across timepoints. Enriched GO terms with false discovery rate (FDR) of less than 0.05 were considered statistically significant and were selected for subsequent downstream analysis.

### Cell-cell communication inference

To analyze intercellular communication network systematically in our snRNA-seq data, normalized data subsets for each respective timepoint (Day 0, 1, 3, and 7) and condition (control vs. plucked), along with cell type labels, were separately submitted for cell-cell communication network analysis using CellChat v2.1.1^57^. Overexpressed signaling genes were identified using Wilcoxon rank sum test. CellChat then inferred statistically and biologically significant ligand-receptor pairs between every pair of cell types. In the main inference analyses, we used the projectData function to smooth the data by projecting the gene expression profiles onto the protein-protein interaction network from STRINGdb^81^. CellChat computed the interaction strength between each ligand-receptor pair, which was quantified by a communication probability value. CellChat employed the law of mass action for calculation of probability value, based on the average expression of each ligand, receptor, and their cofactor(s). Finally, we compared intercellular interactions across conditions by merging CellChat objects.

### Quantitative comparison of cell-cell communication networks

To comprehensively identify signaling alterations between conditions (control vs. plucked) and across timepoints (Day 0, 1, 3, and 7), we employed a top-down approach as previously reported^82^. Briefly, we first investigated global signaling changes at the level of individual cell populations using differential incoming and outgoing interaction strength associated with each cell type, and then executed a refined analysis of perturbed signaling pathways and underlying ligand-receptor pairs.

In the global comparative analysis, we applied network centrality analysis as previously reported^57^. In brief, we used the netAnalysis_computeCentrality function in CellChat to compute the likelihood of each cell type being signaling sources and targets, by summarizing each cell-type-associated outgoing and incoming interaction strengths across all signaling pathways. The netAnalysis_diff_signalingRole_scatter function in CellChat was then performed to compute differential outgoing and incoming centralities of each cell type at each timepoint (plucked centralities – control centralities) and visualize them in a 2-dimensional space for identification of cell types with significant changes in sending or receiving signals between conditions and across timepoints. To better analyze the variations of signaling changes across time, we used the trendline plots to visualize temporal differential outgoing and incoming signaling changes of each cell type for identification of outstanding cell types in each major cell populations (Epithelium, Mesenchyme, and Immune cells). Finally, netVisual_heatmap function, implemented with the ComplexHeatmap^83^ R package, was used to visualize differential cell-cell communication networks among all cell types. This aims to identify outstanding sender-reciver cell pairs at each timepoint. In the heatmap, red and grey colors indicate increased and decreased signaling in plucked, compared to control, respectively.

In the refined analysis at signaling pathway level, we first used the rankNet function in CellChat to identify signaling pathway changes between plucked and control in each outstanding sender-receiver cell pair we identified in the previous cell-level analysis. This aims to compare the communication probabilities of shared signaling pathways between conditions in each sender-receiver pair. The computeCommuProbPathway function in CellChat was used to calculate the communication probability of a given signaling pathway. To identify ligand-receptor pairs that differed significantly between conditions, differential expression testing was applied, in combination with the results of cell-cell communication analysis. Briefly, in each cell type, we used the identifyOverExpressedGenes function to perform Wilcoxon rank sum test for identification of genes differentially expressed between conditions. Signaling molecules were identified as differentially expressed if the *p*-values were less than 0.05 and the absolute values of log-scaled 2-fold difference were higher than 0.25.

## DATA AVAILABILITY

The snRNA-seq data generated in this study are deposited and publicly available in the National Center for Biotechnology Information (NCBI) Gene Expression Omnibus (GEO) under accession GSE315304. The reference genome used in this study was the *Maylandia zebra* UMD2a RefSeq assembly, deposited and publicly available in the NCBI BioProject databank under accession code GCF_000238955.4.

## CODE AVAILABILITY

Code used for core analyses in this study is publicly available at https://github.com/Haowen-He/snRNAseq-Analysis-Accelerated-Tooth-Replacement. The processed data for snRNA-seq is available from Zenodo under https://doi.org/10.5281/zenodo.17983684.

## ACKNOWLEDGEMENTS

We thank Harshita Indukuri and Dr. Laxminarayanan Krishnan for their critical roles in the initial development of the Micro-CT pipelines; Nikesh Kumar and Kathryn Leatherbury for their assistance in preprocessing raw snRNA-seq data; and the Georgia Tech Petit Institute Genome Analysis and Molecular Evolution Cores for their integral roles in sample processing and sequencing, respectively. This work was supported by National Institute of Dental and Craniofacial Research (NIDCR) Grant R01DE019637 to J.T.S.

## AUTHOR CONTRIBUTIONS STATEMENT

J.T.S. and T.M. conceived the project. T.M. and J.T.S. performed the experiments. T.M. carried out the statistical analysis. G.W.G. preprocessed the snRNA-seq data. H.H., G.W.G., and A.S. analyzed the snRNA-seq data. H.H., G.W.G., A.S. and T.M. interpreted and discussed the results with significant intellectual contributions from J.T.S. T.M., H.H., and G.W.G. wrote the initial draft of the manuscript. J.T.S. edited the manuscript with input from all authors. J.T.S. supervised the project.

