## Supplemental Figures 1-10 and Supplemental Tables 1-2 for "Cellular basis of accelerated whole-tooth regeneration": Supplementary_Figure_2.pdf

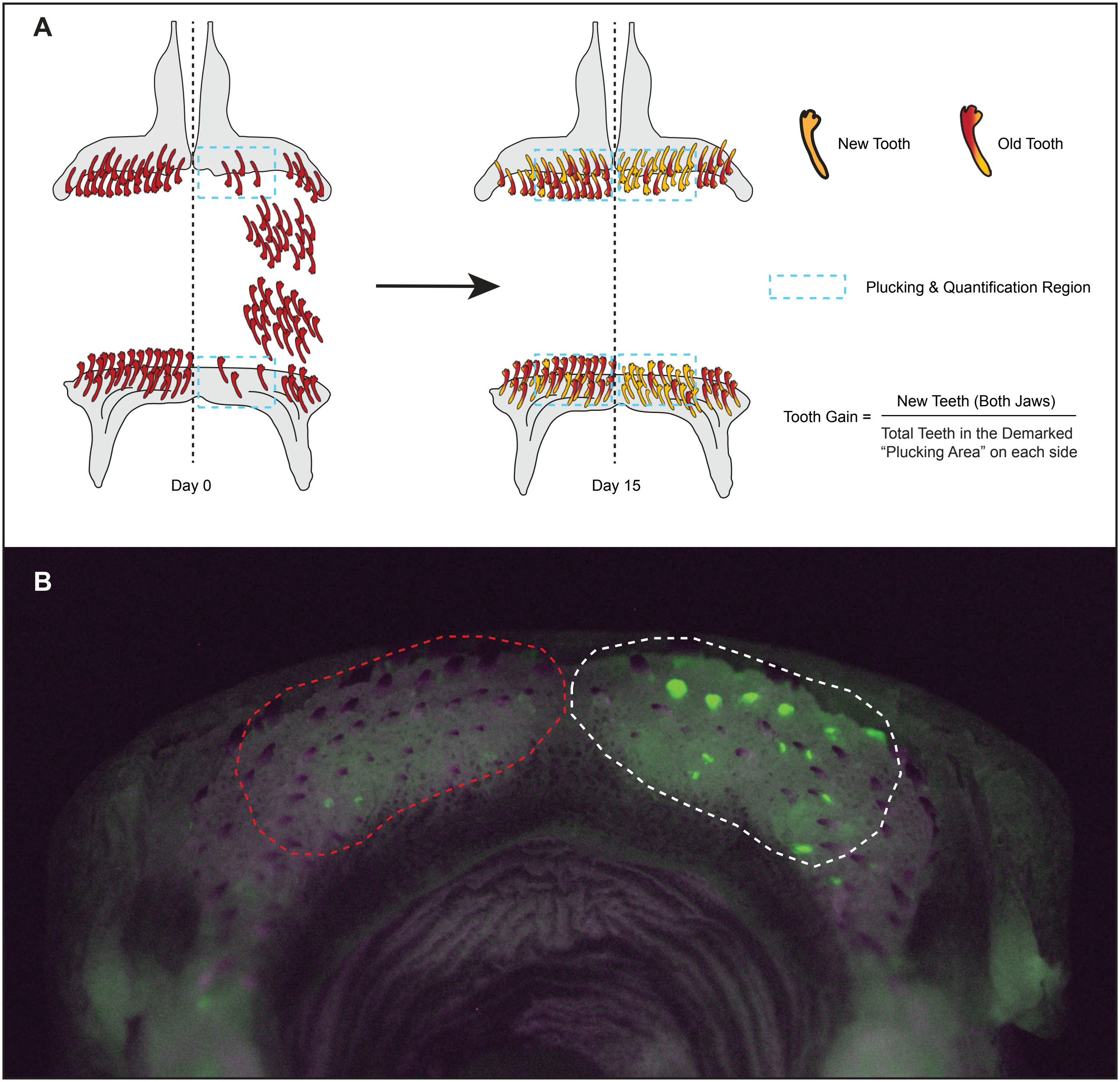


**Figure S2 - Schematic overview of tooth-plucking experiments**. (A) Following Alizarin Red staining, the majority of teeth on the right side of both the upper and lower jaws were removed. Tooth plucking was restricted to the anterio-lateral region of the jaw, as demarcated by the dotted square. Tooth removal was performed under anesthesia as described in the Methods. (B) Representative lower jaw from *Metriaclima estherae* (MZ Red) 15 days post pluck-control experiment. Dotted outlines indicate regions used for quantification: white denotes the plucked side, and red indicates the unmanipulated control side. These regions correspond to the areas used for quantifying newly formed and old teeth on both sides of the jaw. Whole-jaw imaging was acquired using Zeiss Stereo Discovery.V12. Teeth were individually classified and counted using Zeiss Axio Observer Z1 fluorescence.
