## Supplementary figures and images for "Cellular basis of accelerated whole-tooth regeneration"

### Supplementary_Figure_1.jpg

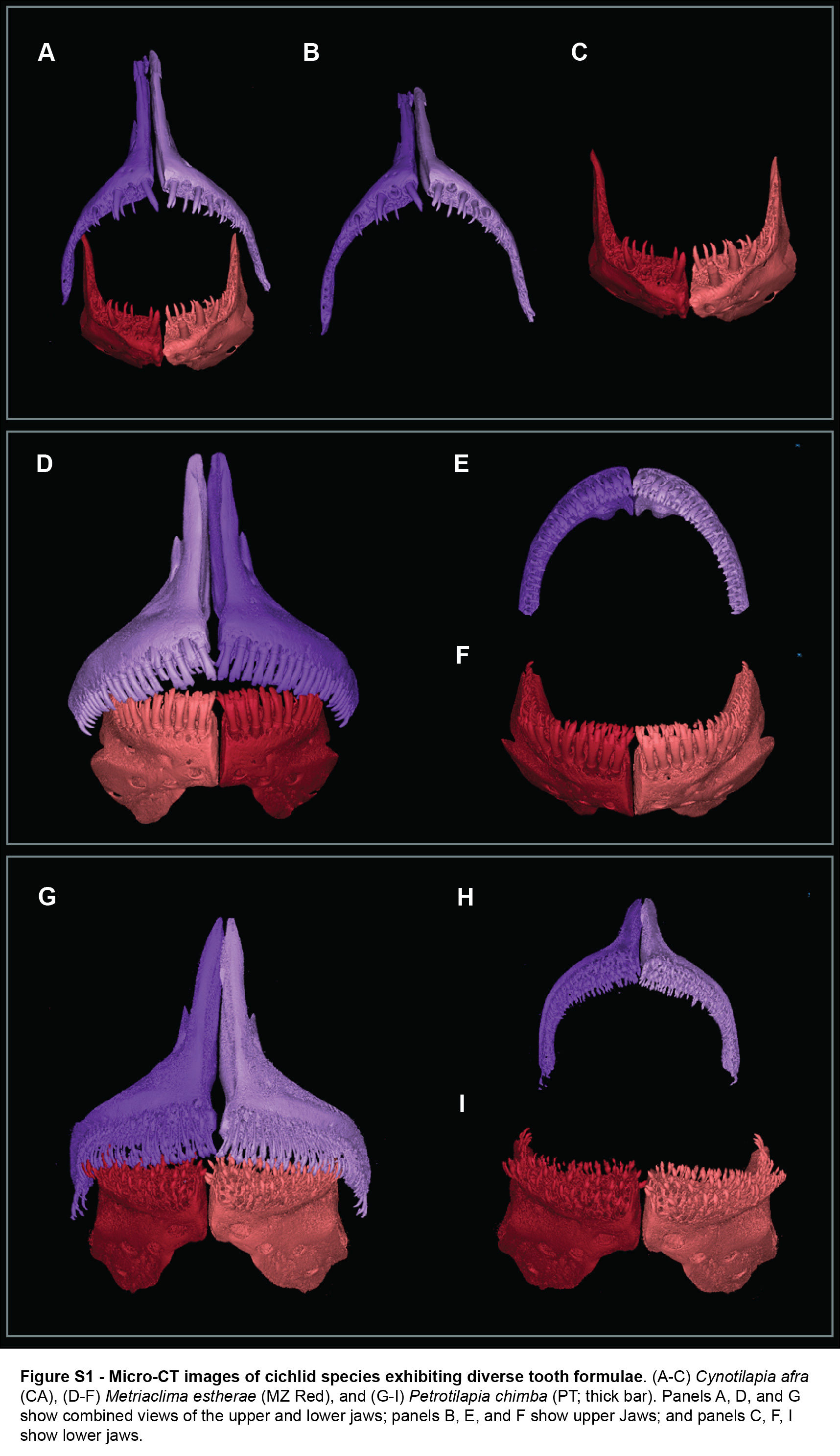

### Supplementary_Figure_3.jpg

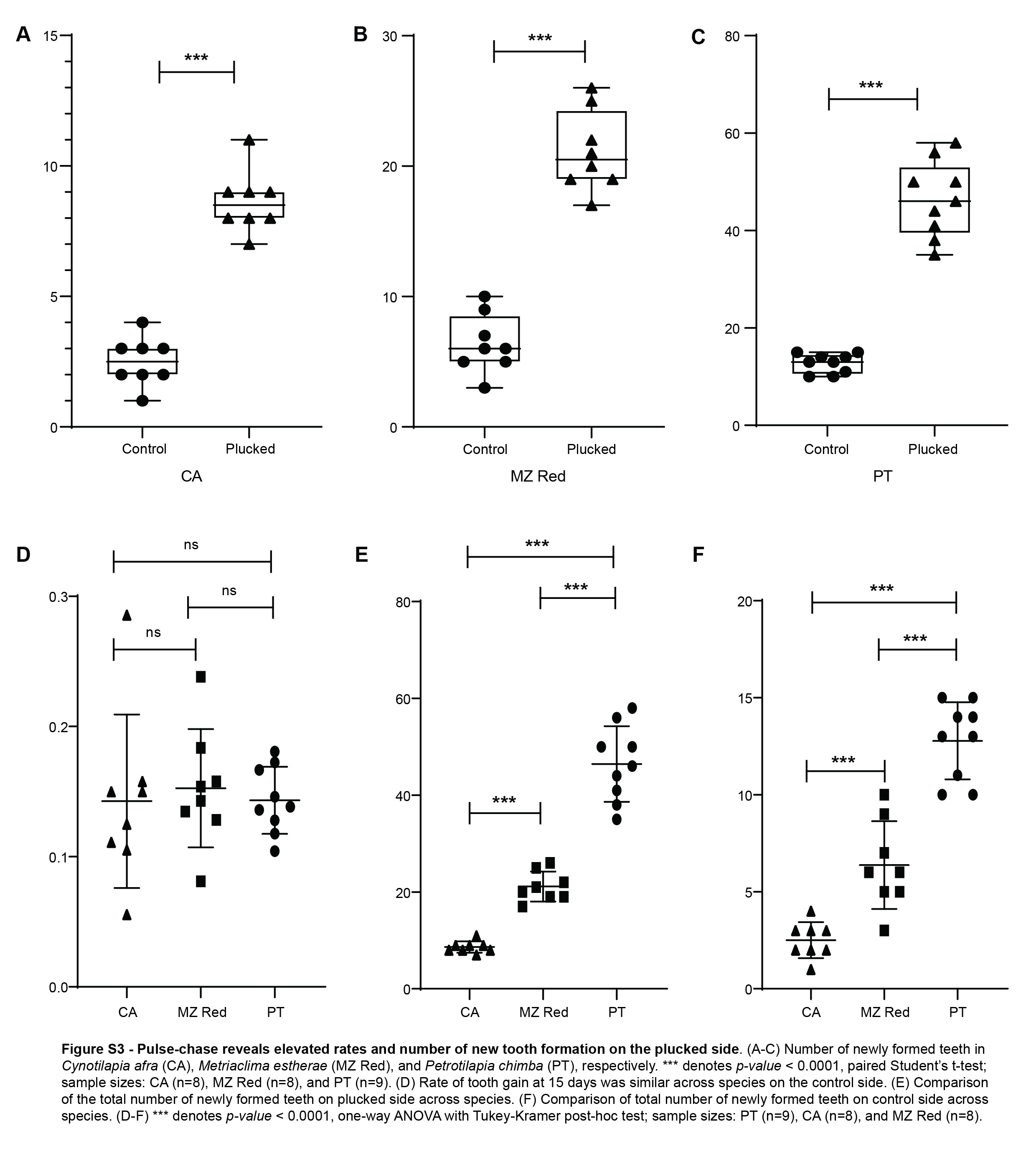

### Supplementary_Figure_4.jpg

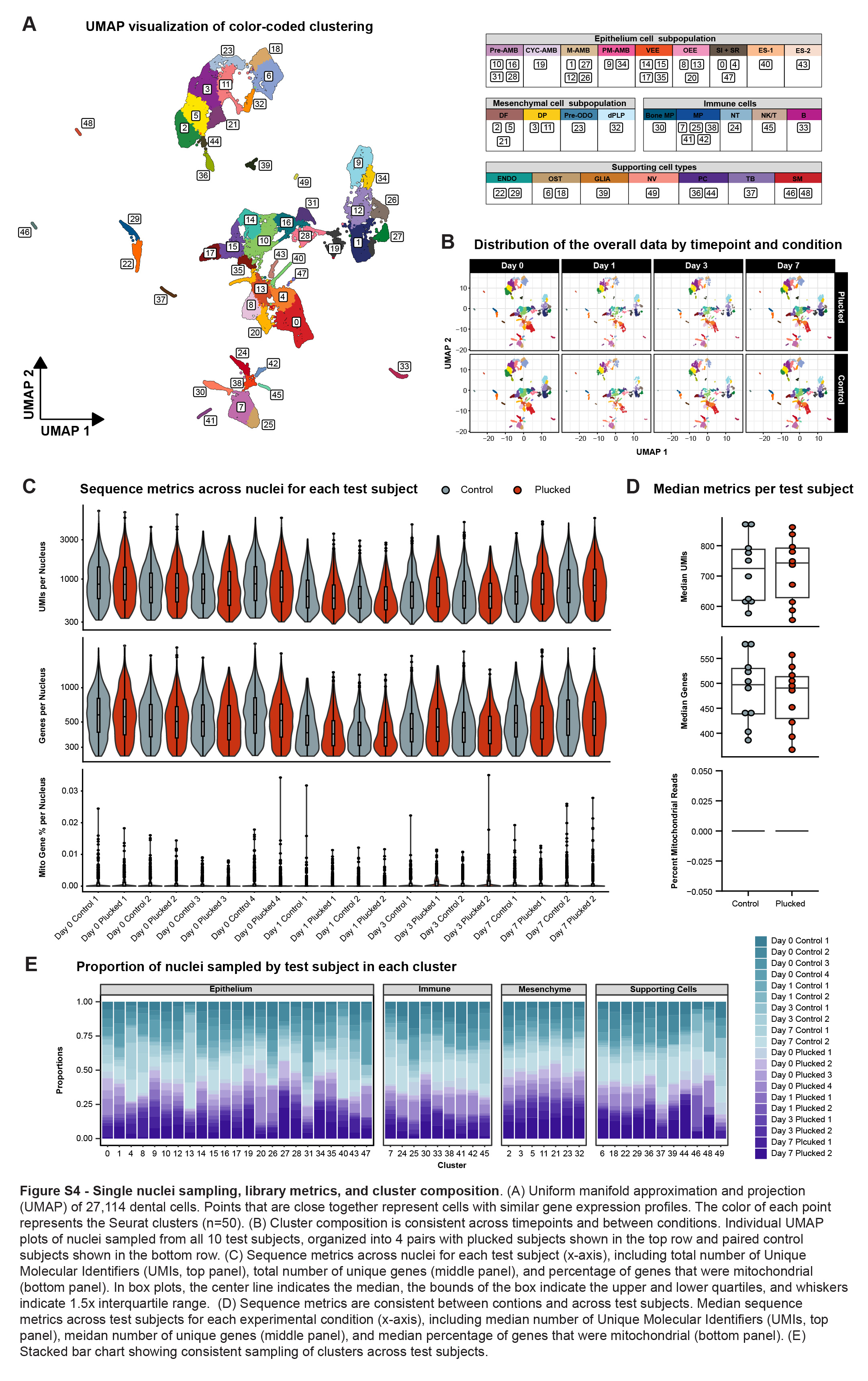

### Supplementary_Figure_5.jpg

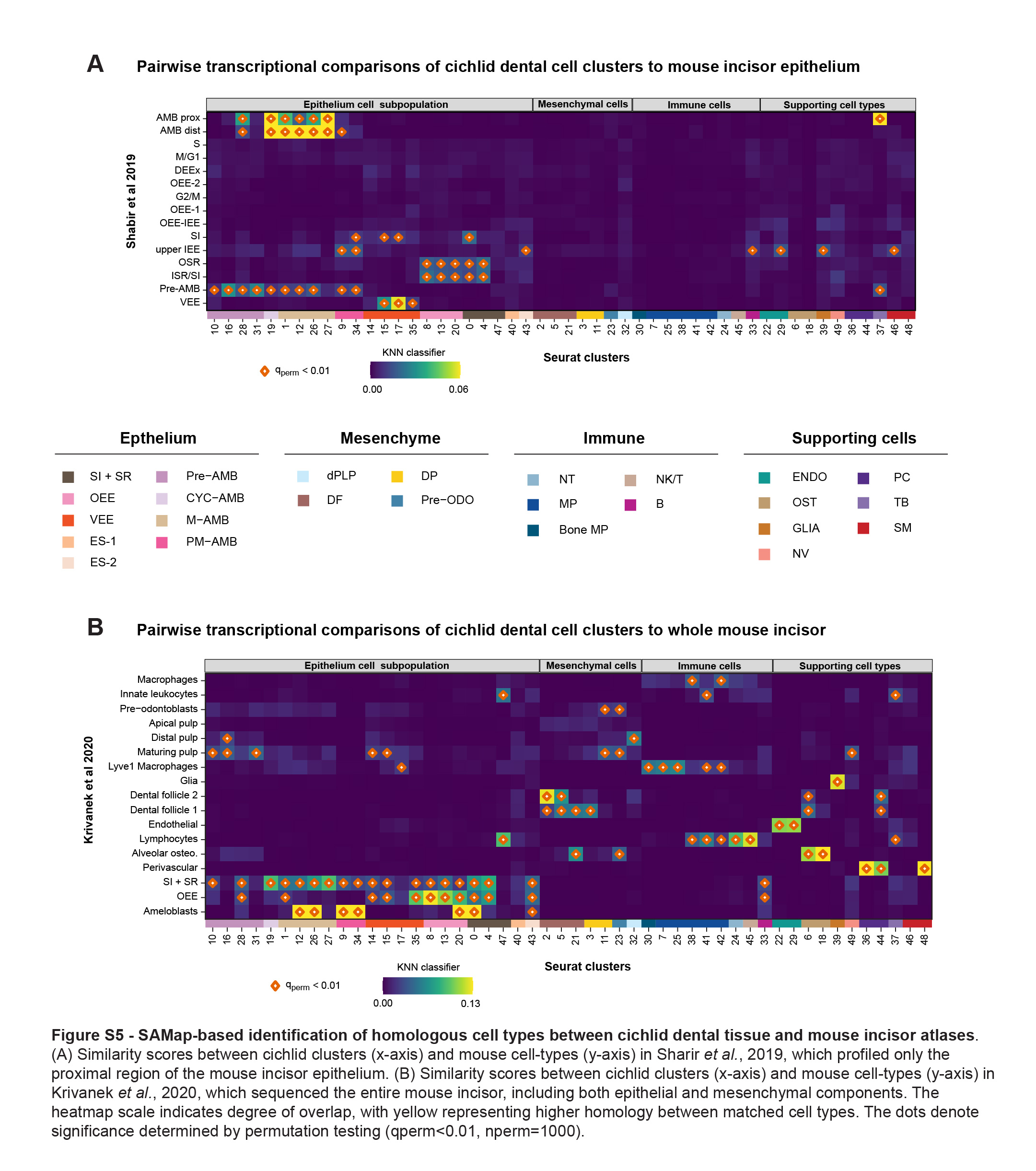

### Supplementary_Figure_6.jpg

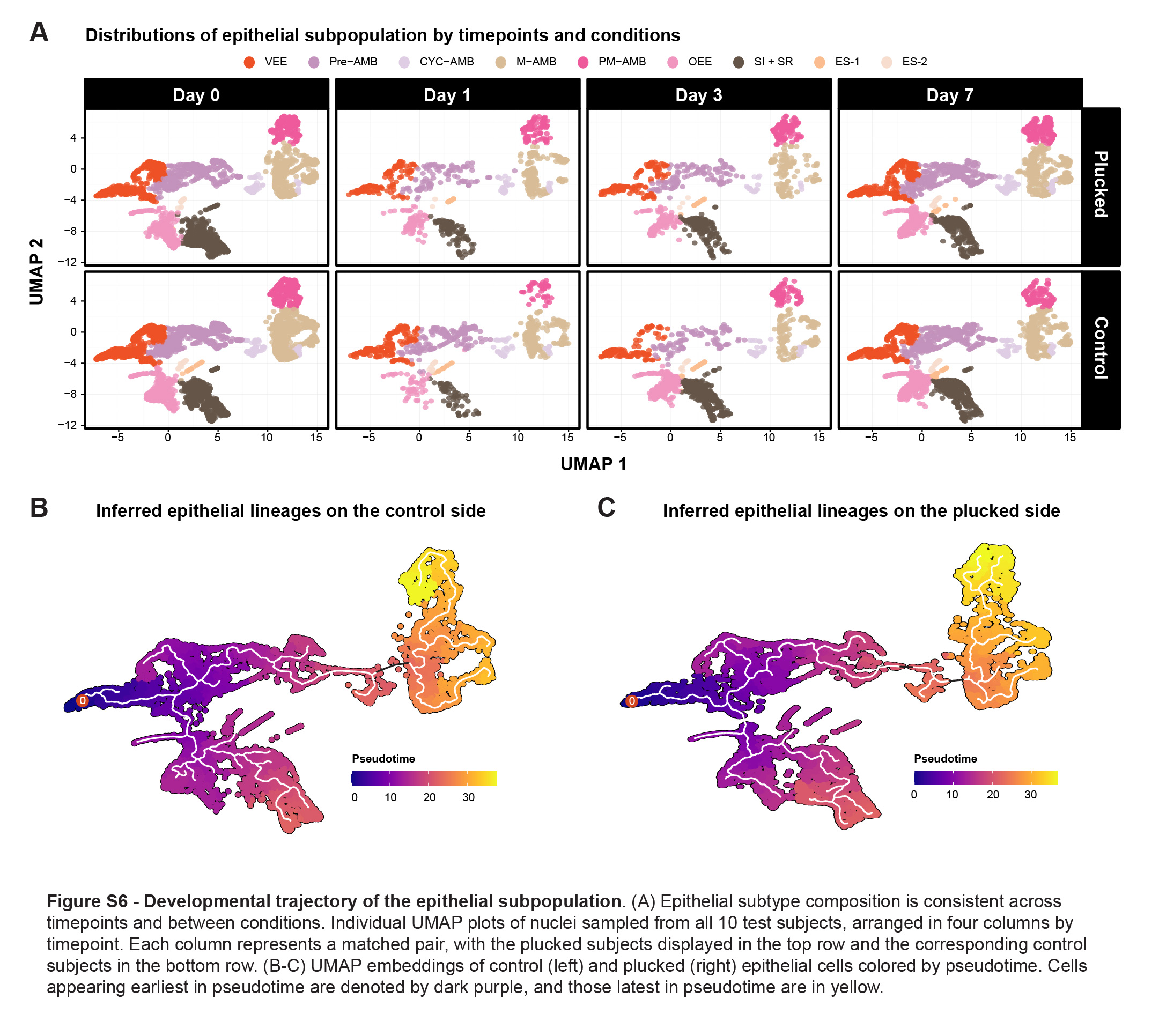

### Supplementary_Figure_8.jpg

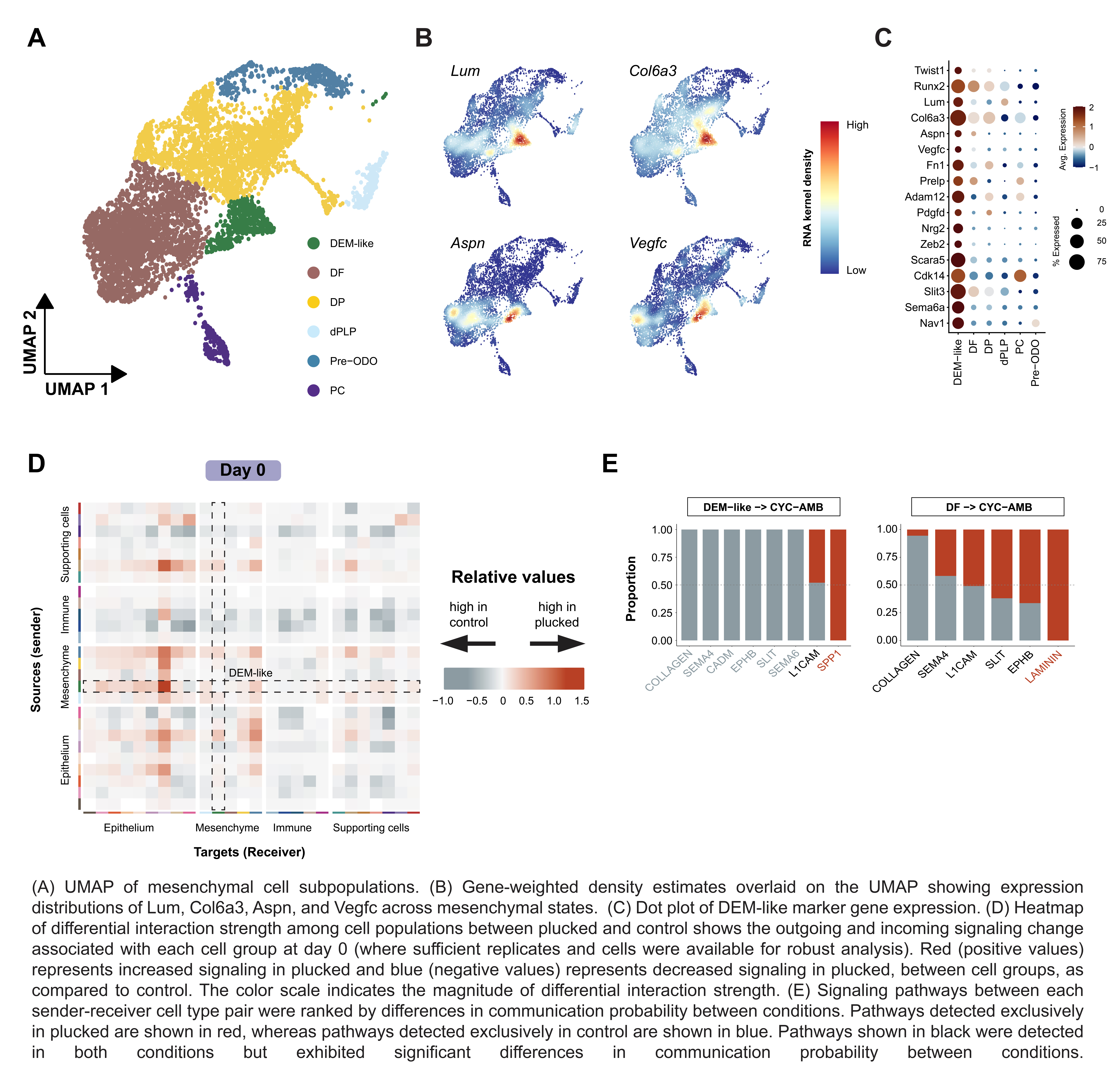

### Supplementary_Figure_9.jpg

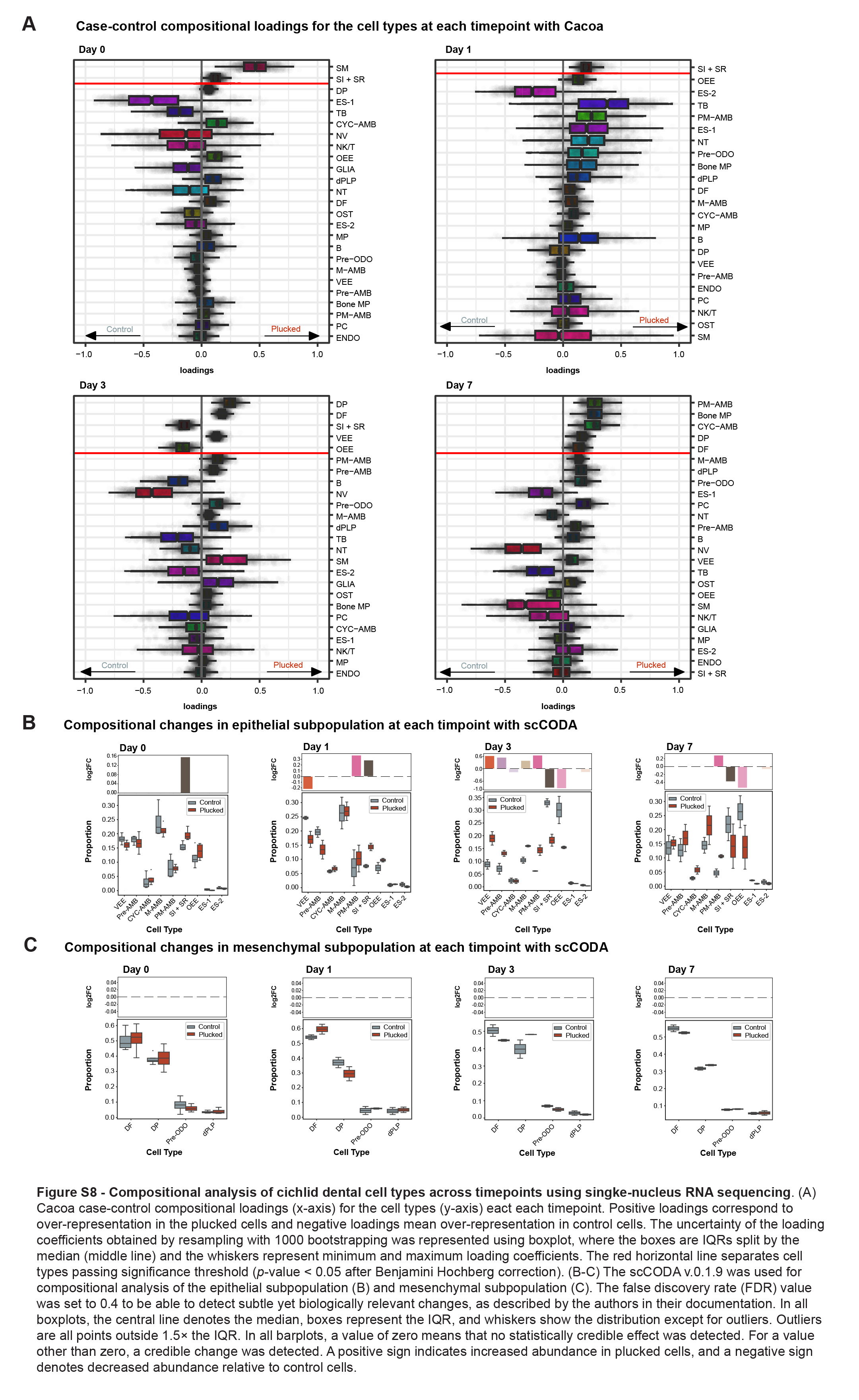

### Supplementary_Figure_10.jpg

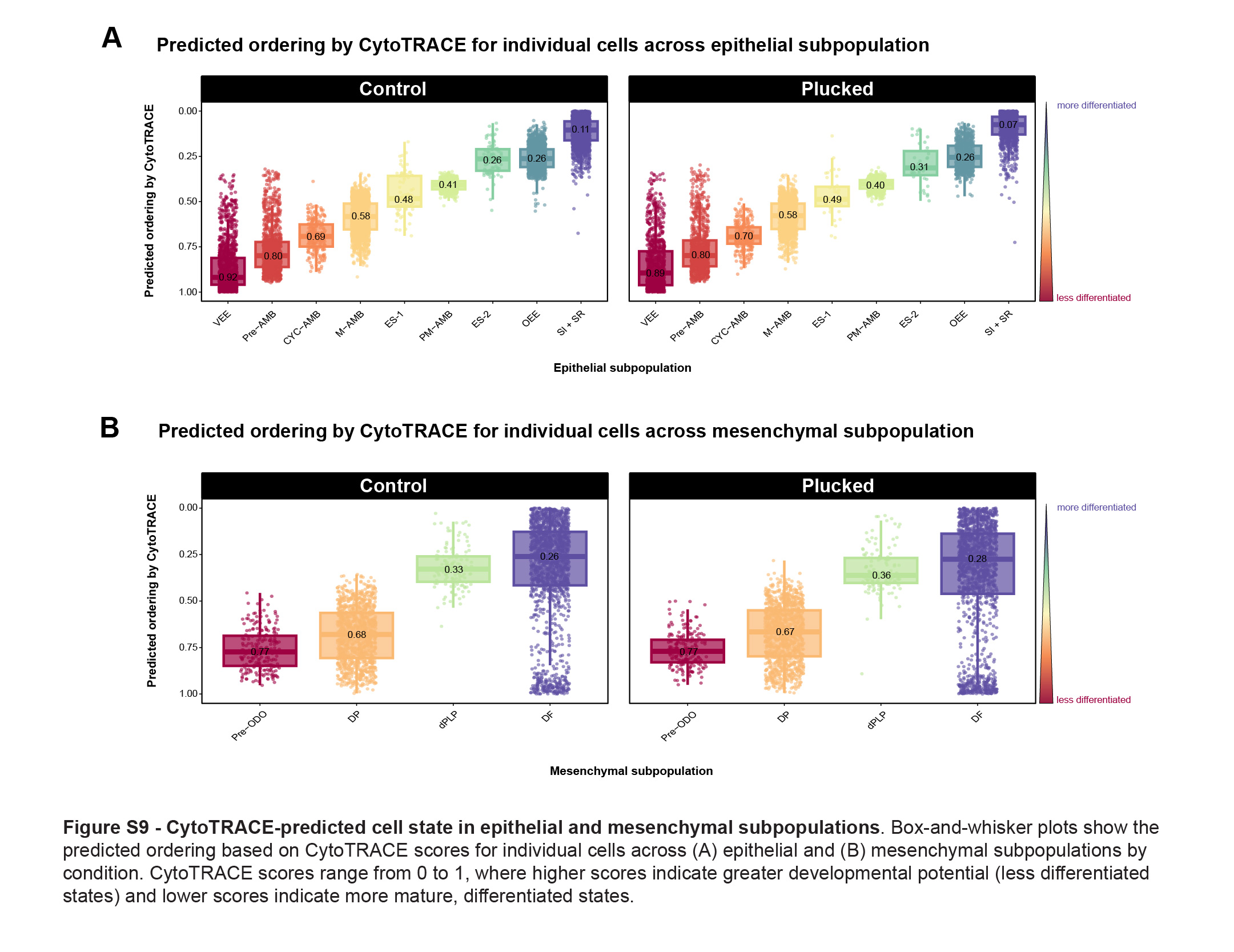
